# Disentangling the biological information encoded in disordered mitochondrial morphology through its rapid elicitation by iCMM

**DOI:** 10.1101/2020.06.22.166165

**Authors:** Takafumi Miyamoto, Hideki Uosaki, Yuhei Mizunoe, Satoi Goto, Daisuke Yamanaka, Masato Masuda, Yosuke Yoneyama, Hideki Nakamura, Naoko Hattori, Yoshinori Takeuchi, Motohiro Sekiya, Takashi Matsuzaka, Fumihiko Hakuno, Shin-Ichiro Takahashi, Naoya Yahagi, Koichi Ito, Hitoshi Shimano

## Abstract

Mitochondrial morphology is dynamically changed in conjunction with spatiotemporal functionality. Although considerable efforts have been made to understand why abnormal mitochondrial morphology occurs in various diseases, the biological significance of mitochondrial morphology in states of health and disease remains to be elucidated owing to technical limitations. In the present study, we developed a novel method, termed inducible Counter Mitochondrial Morphology (iCMM), to purposely manipulate mitochondrial morphological patterns on a minutes timescale, using a chemically inducible dimerization system. Using iCMM, we showed that mitochondrial morphological changes rapidly lead to the characteristic reconstitution of various biological information, which is difficult to investigate by conventional genetic engineering. The manipulation of mitochondrial morphology using iCMM can improve our understanding of the interplay between mitochondrial morphology and cellular functions.

## Introduction

Mitochondria are organelles in eukaryotic cells that have evolved from endosymbiotic α-proteobacteria^1^. They perform numerous roles, and act most prominently as an energy supply machinery that generates the energy currency adenosine triphosphate (ATP) through oxidative phosphorylation^2^. Furthermore, mitochondria are involved in anabolic and catabolic reactions, including, but not limited to, the tricarboxylic acid cycle, β-oxidation of fatty acids, and heme biosynthesis. The products of those reactions are used in various cellular processes, including as building blocks of cellular structure, as substrates in chemical reactions, and as molecular cues in signal transduction networks, indicating that mitochondria are unequivocally indispensable to cells.

As their name implies (Greek mitos = thread; chondrion = granule), mitochondria are highly dynamic organelles that can alter their size, shape, and subcellular distribution through repetitive, coordinated fusion and fission cycles over the course of a few minutes,^3,4^ a phenomenon termed mitochondrial dynamics. Mitochondrial functions have to be coordinated with mitochondrial dynamics, as the latter potentially define the distribution pattern of mitochondrial deliverables, as well as the pattern of inter-organelle communication, in a time-dependent fashion^5,6^. Thus, the spatiotemporal dynamics of mitochondrial morphology are considered an input signal that determines how appropriately mitochondria-involving cell functions are executed under certain conditions. Disturbances in mitochondrial dynamics followed by disruption of mitochondrial morphology occur in various diseases, including cancer and metabolic diseases^7,8^, attesting to the pivotal role of mitochondrial morphology in physiopathology. However, the molecular mechanism underlying the relationship between mitochondrial morphology and cellular functions in health and disease remains unclear.

Over recent decades, considerable efforts have been made to elucidate the biological information encoded in mitochondrial morphology. Generally, mitochondrial morphology was altered either chemically or by manipulating genes involved in mitochondrial morphology^5,9–11^. However, these approaches induced mitochondrial dysfunction and exerted adverse effects on cells because of prolonged arrest of mitochondrial dynamics^12–16^. To overcome this drawback, a tool for inducible, rapid, and specific manipulation of mitochondrial morphology without abrogating mitochondrial functions is required^5^.

Chemically inducible dimerization (CID) systems allow target protein manipulation in a spatiotemporally confined subcellular compartment through a small molecule-induced ternary complex formation of two different proteins^17^. CID systems are applicable to protein-based biocomputing devices for controlling cellular functions^17–20^. Compared to genetic circuit-based biocomputing devices that require a long time (i.e., hours) to execute the logic function, CID systems allow for faster-processing biocomputing devices (seconds to minutes)^18^. Therefore, they are suitable for inducing morphological changes in mitochondria over a short period of time. CID systems have been widely applied in mitochondrial studies, e.g., to promote mitochondria– endoplasmic reticulum (ER) interaction^21^ and to suppress kinase activity in mitochondria^22^. However, a sophisticated platform for manipulating mitochondrial morphology into various patterns using CID systems is lacking.

In the present study, we developed “inducible Counter Mitochondrial Morphology” (“iCMM”), a CID system that utilizes a YES Boolean logic gate, to manipulate mitochondrial morphology in living cells on a minutes timescale. As mitochondrial morphology varies among cell types and conditions, we developed three iCMM systems, each capable of inducing different morphologies. Using iCMM, we determined the type of change occurring immediately following the mitochondrial morphology change, which would otherwise be difficult to study.

## Results

### Development of iCMM

iCMM, a YES Boolean logic gate, consists of an effector (i.e., a designed protein) to perturb mitochondrial morphology and a mitochondria-specific anchor protein that tethers the effector to the mitochondria in the presence of a chemical dimerizer (Fig. 1a). We adopted a rapamycin-based CID system^17^, whereby peptidyl-prolyl cis-trans isomerase FKBP1A (termed FKBP) and the FKBP12-rapamycin binding domain in mammalian/mechanistic target of rapamycin (termed FRB) were served as an effector and an anchor, respectively, using rapamycin as a chemical dimerizer (Fig. 1b).

**Fig. 1:**
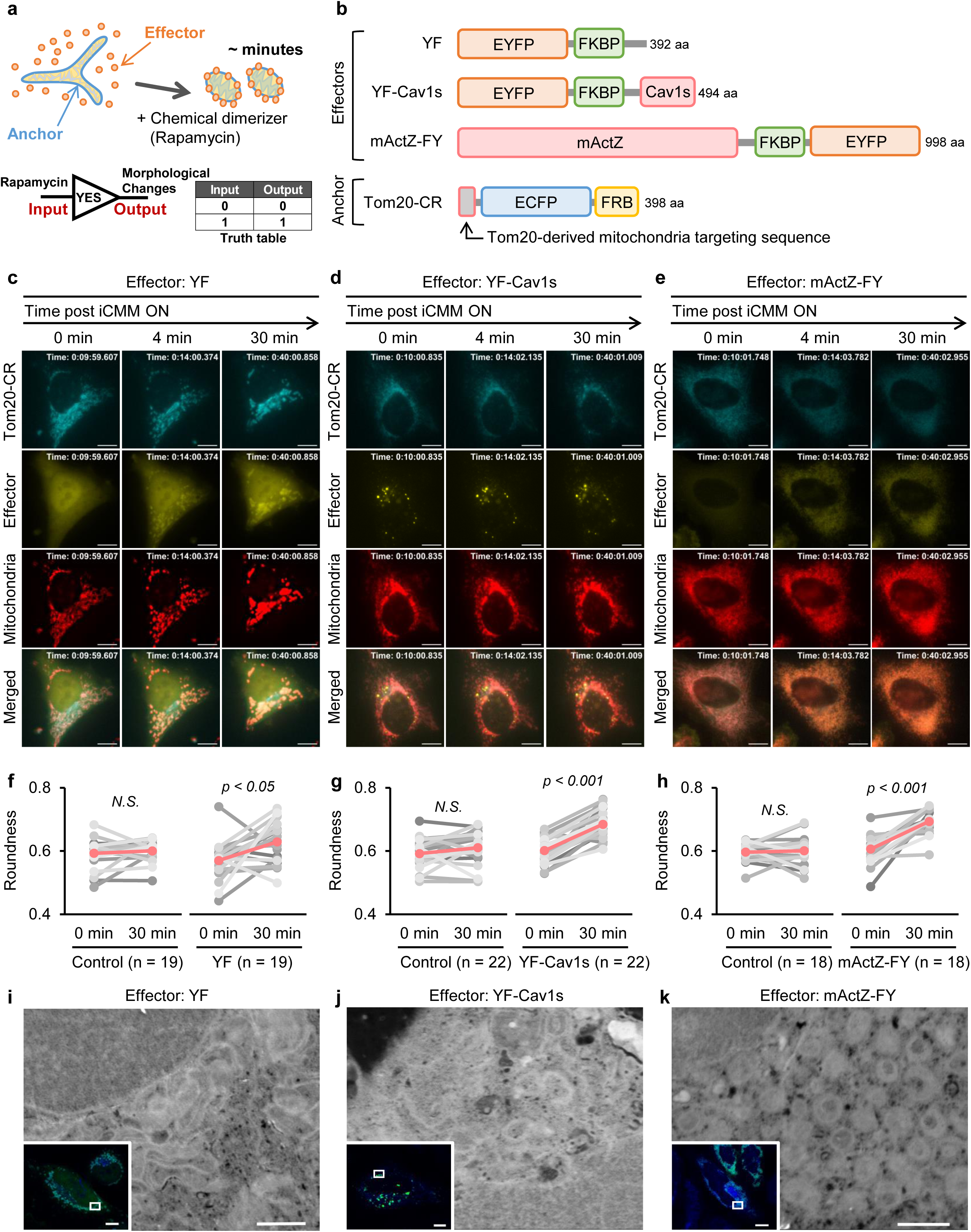
iCMM as a novel CID system for rapid manipulation of mitochondrial morphology in a living cell. **a**, Schematic diagram of the iCMM system. **b**, Structural features of iCMM effectors and anchor. **c–e**, Time-lapse images of HeLa cells expressing Tom20-CR and YF (c), YF-Cav1s (d), or mActZ-FY (e) as iCMM anchor and effectors, respectively. Mitochondria (red) were stained with MitoTracker Red CMXRos; 2 min/frame. Rapamycin was added after image acquisition at frame 6 (indicated as “0 min”). Scale bar: 10 μm. **f–h**, Mitochondrial roundness in HeLa cells expressing indicated iCMM and in surrounding iCMM-negative cells (control) was analyzed before (0 min) and after (30 min) activating iCMM system. Quantification was performed on three independent experiments. Gray: analyzed single cell; Red: average. *p* = paired t-test. *N*.*S*.: statistically nonsignificant. **i–k**, Mitochondrial structures in HeLa cells expressing indicated effectors and Tom20-CR were examined with CLEM after 30 min of iCMM activation. Merged fluorescence images (blue: Tom20-CR, green: indicated effector) and scanning electron microscopy (SEM) images are shown. White boxes in merged fluorescence images indicate areas shown as SEM images. Scale bar: 10 μm (merged fluorescence images), 1 μm (SEM images).

We first used a reported fundamental fusion protein consisting of enhanced yellow fluorescent protein (EYFP) and FKBP (termed YF)^23^ as a functional iCMM effector (Fig. 1b). Although YF has been used as a control effector in various experimental settings^23,24^, we found that once the diffusive YF was substantially translocated to the outer surfaces of mitochondria, which was circumscribed by anchor proteins—which consisted of mitochondrial targeting sequences derived from Tom20, enhanced cyan fluorescent protein (ECFP), and FRB (termed Tom20-CR)—mitochondrial morphology changed from a basal reticular thread-like structure to large punctate structures (Fig. 1c, Supplementary movie 1). This mitochondrial network disruption was confirmed as an increase in the roundness of mitochondria (Fig. 1f). Correlative light and electron microscopy (CLEM) indicated that YF translocation induced mitochondrial crowding, not fusion (Fig. 1i). mYF containing monomeric EYFP^A206K^ (mEYFP) instead of EYFP did not change mitochondrial morphology, although it substantially accumulated in the mitochondrial outer membrane (Supplementary Fig. 1, Supplementary movie 2), suggesting that the inherent homodimerization activity of EYFP^25^ was, at least partially, required for the YF-induced change in mitochondrial morphology. We used mYF as a reference for the functional iCMM effectors in subsequent experiments.

We next focused on caveolin-1, an integral membrane protein that localizes to caveola and multiple interior compartments through vesicle trafficking^26^. As fluorescent protein-fused caveolin-1 exhibits a punctate distribution pattern in the cell^27^, we hypothesized that the stabilization of mitochondria with the punctate structures via the CID system could induce mitochondrial morphology change. For this concept, we constructed a functional iCMM effector consisting of YF and amino acids (aa) 61–178 of caveolin-1 (termed YF-Cav1s, Fig. 1b), which was the smallest fusion protein showing a punctate distribution pattern similar to that of fluorescent protein-fused caveolin-1 (data not shown). Following the addition of rapamycin, mitochondrial morphology changed from a network structure to punctate structures of various sizes in cells expressing YF-Cav1s and Tom20-CR (Fig. 1d, g, and j; Supplementary movie 3).

Lastly, we optimized a previously designed interspecies fusion protein consisting of aa 30–262 of *Listeria monocytogenes* Actin assembly-inducing protein (ActA), codon-optimized for usage in mammalian cells^24^, and aa 2–380 of human zyxin (termed mActZ), and being structurally and functionally similar to full-length ActA^28^. As ActA converts actin polymerization into a motile force^29^, we hypothesized that the mechanical force generated by actin polymerization could alter mitochondrial morphology. mActZ was fused with FKBP and EYFP (termed mActZ-FY, Fig. 1b) to produce the functional iCMM effector. Similar to ActuAtor, a codon-optimized ActA functionally homologous to mActZ-FY^24^, cytosolic mActZ-FY recruitment to mitochondria by rapamycin treatment obviously transformed mitochondrial morphology to small, dot-like structures of almost equal size, in cells expressing mActZ-FY and Tom20-CR (Fig. 1e, h, and k, Supplementary movie 4).

Together, these results indicate that each functional iCMM effector could alter mitochondrial morphology in different ways on a minutes timescale.

### iCMM specifically alters mitochondrial morphology in target cells

In addition to the timescale, we confirmed the specificity of iCMM; none of the functional iCMM effectors developed in the present study changed mitochondrial morphology prior to activation of the iCMM system, as judged from the average area and number of mitochondria in the cells (Supplementary Fig. 2). Notably, the average mitochondrial area and number could not be distinguished in cells expressing iCMM with YF and Tom20-CR before and after operating the iCMM system (Supplementary Fig. 2e, l). This is because YF induced mitochondrial assemblage, but not intense fragmentation, as observed for YF-Cav1s and mActZ-FY (Fig. 1). We confirmed that iCMM specifically induced mitochondrial morphology changes, without affecting the morphology of other organelles during a 120-min observation window (Supplementary Fig. 3).

One of the strengths of the CID system is its versatility^17,30^. To determine versatility, the iCMM systems were transduced into Hep G2 human liver cancer cell line and U-2 OS human osteosarcoma cell line. Each functional iCMM effector induced a characteristic mitochondrial morphology change in both cell lines, on a similar timescale (Supplementary Fig. 4).

### iCMM alters mitochondrial morphology without loss of mitochondrial membrane potential

The mitochondrial membrane potential (ΔΨm), which results from redox transformations, plays decisive roles in various cellular functions, including ATP synthesis and innate immune responses^31–33^. Remarkably, ΔΨm is also critical in determining mitochondrial morphology^34^. To resolve the biological significance of mitochondrial morphology in various cellular functions, devices capable of inducing mitochondrial morphology changes at specific times, without altering ΔΨm in the target living cells, are needed. Therefore, we examined the effect of iCMM on ΔΨm. Regardless of the effector used, tetramethylrhodamine ethyl ester (TMRE) staining revealed that no change in ΔΨm occurred before or after mitochondrial morphology change induction by iCMM, during a 120-min time window (Fig. 2a–d). We further examined whether mitochondrial morphology changes induced by iCMM altered the susceptibility of mitochondria to FCCP, an ionophore uncoupler of oxidative phosphorylation (Supplementary Fig. 5a). Time-lapse imaging revealed that FCCP treatment resulted in loss of ΔΨm, irrespective of mitochondrial morphology (Fig. 2e–h; Supplementary Fig. 5b). Together, these results indicated that iCMM altered mitochondrial morphology without affecting ΔΨm.

**Fig. 2:**
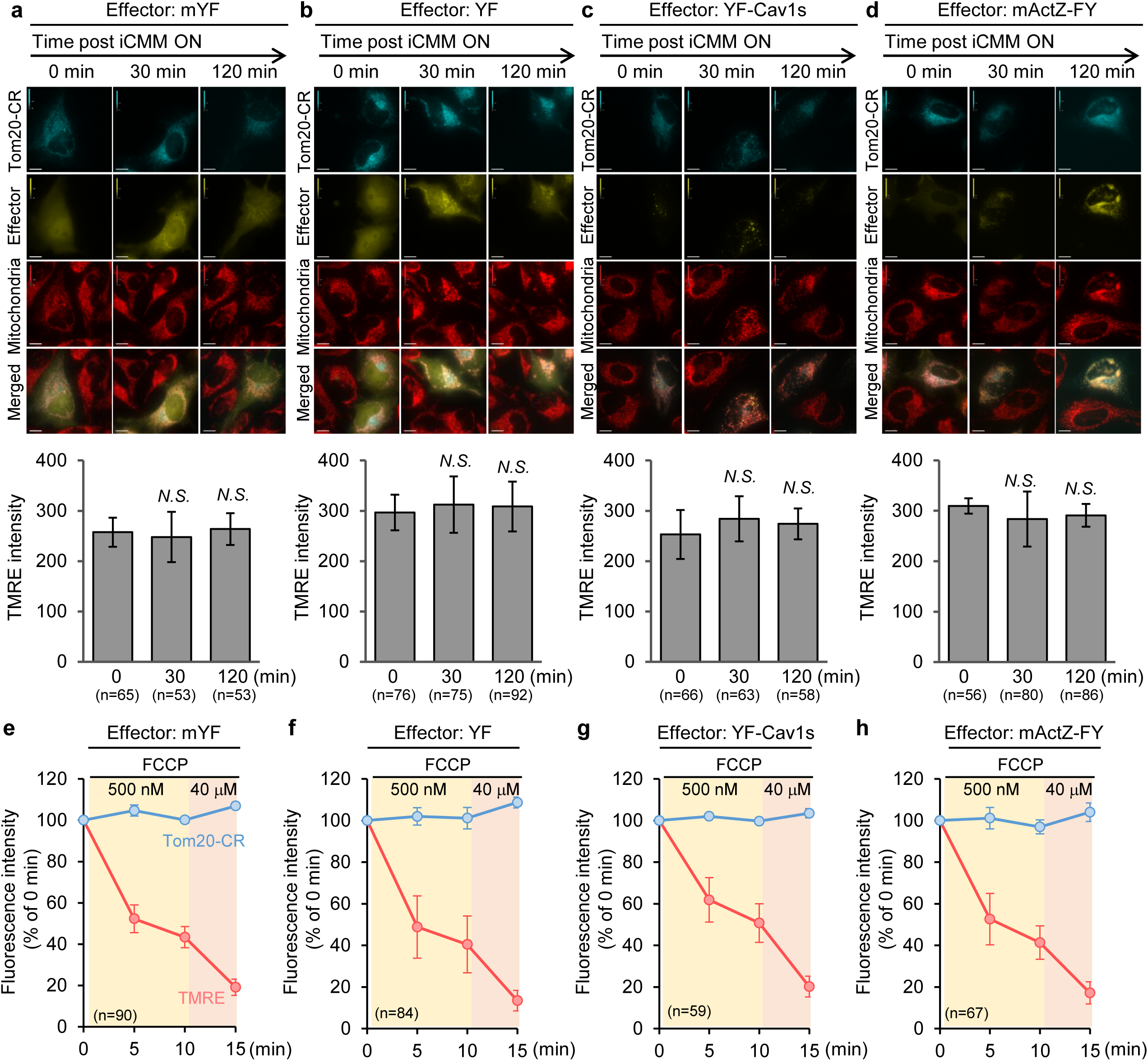
iCMM alters mitochondrial morphology without loss of ΔΨm. **a–d**, ΔΨm in HeLa cells expressing the indicated iCMM system was measured at each indicated time point after activating iCMM system. Mitochondria (red) were stained with TMRE. Representative images at each time point are shown. Quantification was performed on three independent experiments. All data are presented as the mean ± standard deviation. *N*.*S*.: statistically nonsignificant (Student’s t-test). **e–h**, Time-lapse images of ΔΨm in HeLa cells expressing Tom20-CR and indicated iCMM effector are shown; 5 min/frame. FCCP was added after image acquisition at frame 1. All data are presented as the mean ± standard deviation obtained from three independent experiments. Quantification is shown in Supplementary Fig. 5.

### Disruption of mitochondrial morphology by mActZ, but not other effectors, suppresses oxygen consumption by mitochondria

In mammalian cells, mitochondria are responsible for the majority of cellular oxygen consumption to generate ATP. To maintain bioenergetic activity in this process, mitochondria repeatedly go through fusion and fission cycles, ^14,35,36^. To examine the effect of iCMM-induced mitochondrial morphology disruption on the mitochondrial respiration capacity, we established cell lines stably expressing iCMM. Compared to control cells that stably expressed the control iCMM effector mYF and Tom20-CR (termed iCMM^*mYF*^ cells), no noticeable differences in cell proliferation and cell morphology were observed in cells stably expressing Tom20-CR and a functional iCMM effector (YF, YF-Cav1s, or mActZ-FY) (termed iCMM^*YF*^, iCMM^*Cav1s*^, and iCMM^*mActZ*^, respectively) (Supplementary Fig. 6a, b). As expected, iCMM system activation had similar effects on mitochondrial morphology in the stable cell lines as in cells that transiently expressed iCMM (Supplementary Fig. 6c–f).

We next examined the relationship between mitochondrial morphology and mitochondrial respiration activity by measuring the oxygen consumption rate (OCR). Before OCR measurement, cells were treated with rapamycin to induce mitochondrial morphology change or with DMSO as a control for 30 min. Although rapamycin can affect the OCR under certain conditions^37,38^, no significant difference in the OCR was observed in iCMM^*mYF*^ cells treated with DMSO or rapamycin, indicating that rapamycin, at least under the current conditions, did not affect the OCR (Fig. 3a). Similarly, disruption of the mitochondrial network by YF and YF-Cav1s did not affect the OCR (Fig. 3b, c). In contrast, rapamycin-treated iCMM^*mActZ*^ cells exhibited had a lower OCR than did control cells (Fig. 3d). Prolonged (2 h) disruption of the mitochondrial network resulted in similar phenotypes (Supplementary Fig. 7). However, the maximal and non-mitochondrial OCR in iCMM^*mActZ*^ cells varied depending on the time elapsed after mitochondrial morphology change occurrence (Fig. 3d and Supplementary Fig. 7d), implying that maximal and non-mitochondrial respiration might be reorganized to adapt to the lowered mitochondrial respiration. Although mitochondrial ATP production activity was decreased in iCMM^*mActZ*^ cells, similar to the other cell lines, the overall ATP level in the cells remained unchanged, as ATP production in proliferating cells is attributed to glycolysis^39^ (Fig. 3e–h). Together, these results suggested that the OCR fluctuated according to the mitochondrial morphology.

**Fig. 3:**
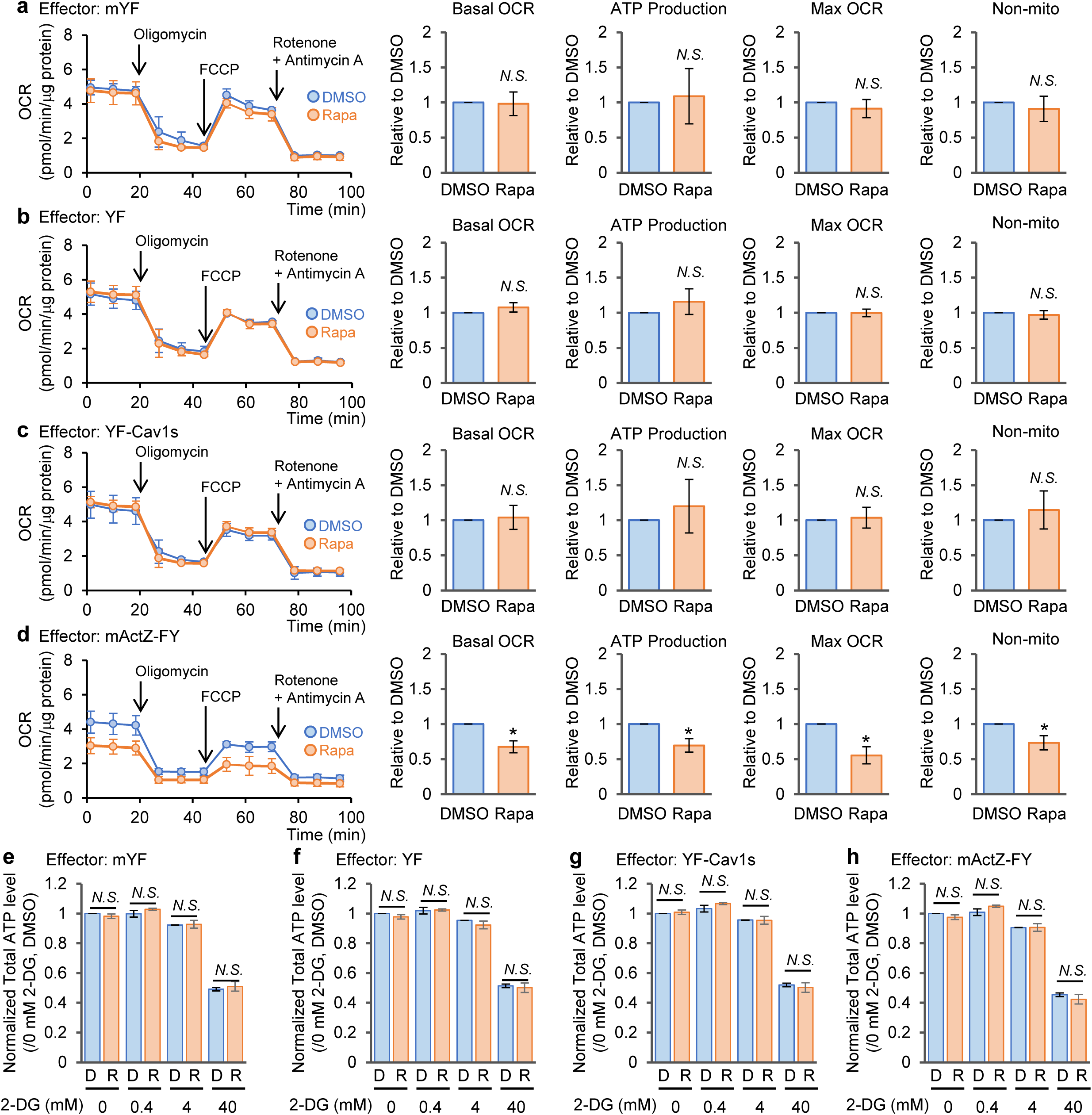
mActZ-FY-induced disruption of mitochondrial morphology reduces the OCR. **a–d**, OCR in iCMM^*mYF*^ (a), iCMM^*YF*^ (b), iCMM^*Cav1s*^ (c), and iCMM^*mActZ*^ (d) are shown. Before OCR measurement, all cell lines were treated with DMSO or rapamycin for 30 min. Quantification was performed on three independent experiments. All data are presented as the mean ± standard deviation. \**P* < 0.05, *N*.*S*.: statistically nonsignificant (Paired t-test). **e–h**, Intracellular ATP level in iCMM^*mYF*^ (e), iCMM^*YF*^ (f), iCMM^*Cav1s*^ (g), and iCMM^*mActZ*^ (h) are shown. Each cell line was treated with DMSO (D) or rapamycin (R) for 1 h, followed by 2-DG treatment for 4 h. Quantification was performed on three independent experiments. All data are presented as the mean ± standard deviation. *N*.*S*.: statistically nonsignificant (Student’s t-test).

### Mitochondrial morphology affects parkin accumulation in mitochondria

Mitophagy, the specific autophagic degradation of mitochondria, is at the core of mitochondrial quality control^40^. During this process, the E3 ubiquitin ligase parkin selectively translocates to dysfunctional mitochondria with low ΔΨm^40^. Although mitochondria that potentially are a mitophagy substrate usually are fragmented, mitochondrial fragmentation *per se* is not sufficient to cause parkin recruitment^41^. These findings were made after inducing mitochondrial morphology change for an extended period; therefore, we reexamined the interplay between mitochondrial morphology and parkin translocation using iCMM, which can alter mitochondrial morphology on a minutes timescale. Cells transiently expressing iCMM were treated with rapamycin for 2 h to examine whether the mitochondrial morphology change *per se* would cause parkin translocation (Supplementary Fig. 8). Parkin recruitment was observed in a small proportion of mitochondria whose morphology was altered by YF-Cav1s and mActZ-FY (Fig. 4a). Next, we tested whether mitochondrial morphology affected FCCP-induced parkin translocation, excluding cells that showed parkin translocation to mitochondria before FCCP treatment. (Supplementary Fig. 8). Compared to control cells that expressed mYF, mitochondrial fragmentation induced by YF-Cav1s and mActZ-FY did not promote parkin translocation to mitochondria in the presence of FCCP (Fig. 4b). In contrast, mitochondrial crowding induced by YF suppressed FCCP-induced parkin translocation (Fig. 4b). Time-lapse imaging revealed that the speed of parkin translocation did not differ between the various mitochondrial morphologies induced by the iCMM systems (Fig. 4c, d). Considering that FCCP treatment decreased the ΔΨm in an mitochondrial morphology-independent manner (Fig. 2e–h), these results implied that parkin recruitment to depolarized mitochondria might be, at least partially, regulated by mitochondrial morphology-related mechanisms.

**Fig. 4:**
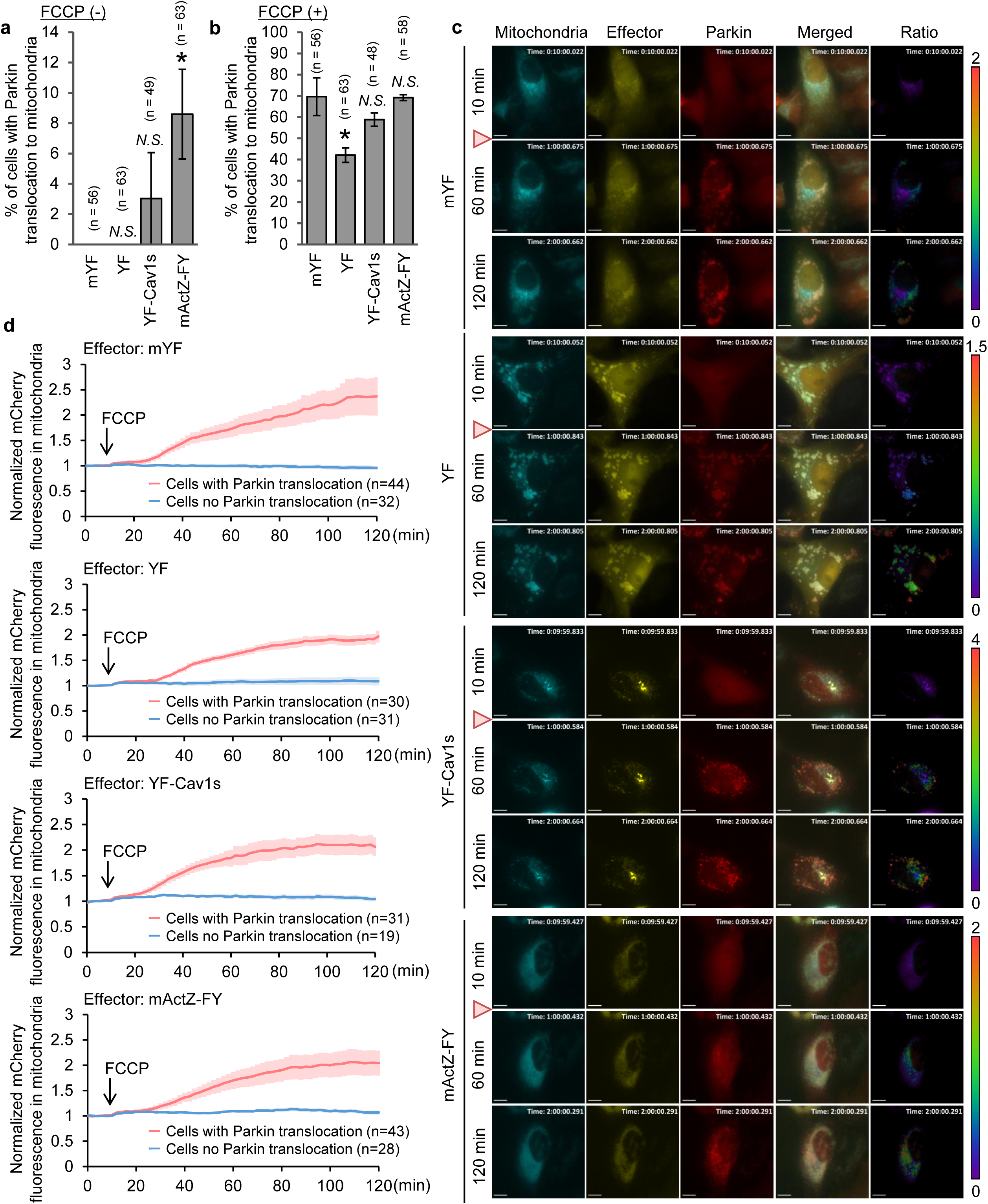
Translocation efficiency of Parkin to mitochondria in the presence of FCCP is partially regulated by mitochondrial morphology. **a**, HeLa cells transiently expressing Tom20-CR, the indicated iCMM effector, and mCh-Parkin were treated with rapamycin for 2 h. Quantification was performed in three independent experiments. All data are presented as the mean ± standard deviation. \**P* < 0.05, *N*.*S*.: statistically nonsignificant (Student’s t-test). **b**, HeLa cells from (a) not showing Parkin translocation to the mitochondria were treated with 10 μM FCCP for 110 min, then cells showing Parkin translocation to the mitochondria were counted. Quantification was performed in three independent experiments. All data are presented as mean ± standard deviation. \**P* < 0.05, *N*.*S*.: statistically nonsignificant (Student’s t-test). **c**, HeLa cells not showing Parkin translocation to the mitochondria were subjected to time-lapse imaging; 1 min/frame. FCCP (10 μM) was added after image acquisition at frame 11 (red triangle). Representative images are shown. Scale bar: 10 μm. Mitochondria: Tom20-CR, Effector: indicated iCMM effector, Parkin: mCh-Parkin. Ratio = Parkin/mitochondria. **d**, Time-lapse images of Parkin in mitochondria in the cells from (c) are shown. All data are presented as the mean ± standard deviation.

### Mitochondrial morphology change does not render cells susceptible to staurosporine-induced apoptosis

During apoptosis, large intracellular structures reorganize to prepare for cell death^42^. One hallmark of this reorganization is mitochondrial fragmentation, caused by an imbalance in the mitochondrial fusion-fission cycles^43,44^. It should be noted that mitochondrial fragmentation could determine susceptibility to apoptosis-inducible input^45–47^. To examine whether iCMM-induced mitochondrial morphology change affected susceptibility to apoptosis induction, cells stably expressing iCMM were treated with staurosporine (STS), an apoptosis inducer. Disrupted mitochondrial morphology did not promote STS-induced caspase-3 activation (Fig. 5a). Notably, rapamycin failed to attenuate mTORC1 activity in iCMM^*mActZ*^ cells, whereas STS-induced suppression of mTORC1 activity^48^ was similar to that in other cells (Fig. 5a). Torin 1, an mTOR inhibitor, completely suppressed mTORC1 activity (Supplementary Fig. 9), indicating that the mActZ-FY-induced mitochondrial morphology change attenuated the effect of rapamycin on mTORC1 signaling. Nuclear morphology analysis in cells stably or transiently expressing the iCMM systems supported that mitochondrial morphology was not involved in the sensitivity to STS-induced apoptosis (Fig. 5b, c). Collectively, these results suggested that changes in intracellular information accompanied with changes in mitochondrial morphology, rather than mitochondrial morphology *per se*, played a pivotal role in determining susceptibility to STS-induced apoptosis during the observation window.

**Fig. 5:**
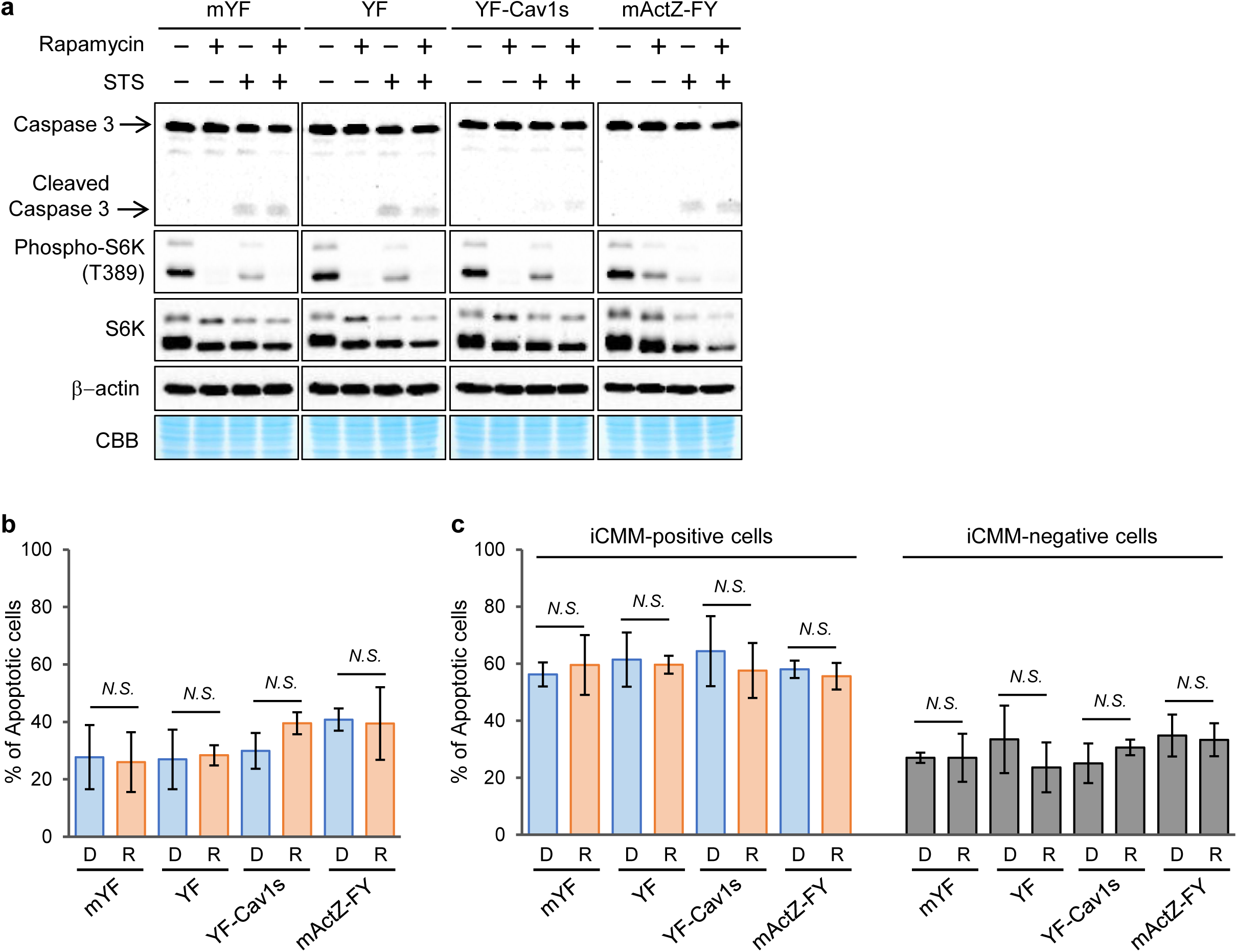
mitochondrial morphology does not affect STS-induced apoptosis. **a**, HeLa cells stably expressing Tom20-CR and the indicated iCMM effector were treated with DMSO or rapamycin for 1 h, followed by treatment with 266 nM STS for 6 h. The cells were then subjected to western blot analysis. **b**, HeLa cells stably expressing Tom20-CR and the indicated iCMM effector were treated as in (a). Subsequently, normal and apoptotic nuclei in the cells were counted. mYF-DMSO: n = 775, mYF-Rapa: n = 974, YF-DMSO: n = 727, YF-Rapa: n = 695, Cav1s-DMSO: n = 864, Cav1s-Rapa: n = 684, mActZ-DMSO: n = 545, mActZ-Rapa: n = 568. Quantification was performed in three independent experiments. All data are presented as the mean ± standard deviation. *N*.*S*.: statistically nonsignificant (Student’s t-test). **c**, HeLa cells transiently expressing Tom20-CR and the indicated iCMM effector were treated as in (a). Subsequently, normal and apoptotic nuclei in cells expressing iCMM (iCMM-positive cells) or peripherally not expressing iCMM (iCMM-negative cells) were counted. For iCMM-positive cells, the following cell numbers were determined: mYF-DMSO: n = 223, mYF-Rapa: n = 211, YF-DMSO: n = 222, YF-Rapa: n = 198, Cav1s-DMSO: n = 194, Cav1s-Rapa: n = 261, mActZ-DMSO: n = 190, mActZ-Rapa: n = 208. For iCMM-negative cells, the following cell numbers were determined: mYF-DMSO: n = 1002, mYF-Rapa: n = 544, YF-DMSO: n = 794, YF-Rapa: n = 735, Cav1s-DMSO: n = 967, Cav1s-Rapa: n = 995, mActZ-DMSO: n = 851, mActZ-Rapa: n = 810. Quantification was performed in three independent experiments. All data are presented as the mean ± standard deviation. *N*.*S*.: statistically nonsignificant (Student’s t-test). D: DMSO, R: Rapamycin.

### Rapid mitochondrial morphology change induced by iCMM alters the transcriptome

While most mitochondrial morphology changes are triggered by changes in the cell milieu, a comprehensive understanding of how mitochondrial morphology changes alter intracellular information is lacking. Although we developed the iCMM system to disentangle physiopathological information embedded in mitochondrial morphology, it is also applicable to the manipulation of intracellular information through mitochondrial morphology reorganization. To elucidate the effects of the iCMM systems on the gene transcription profile, HeLa cells stably expressing iCMM were subjected to transcriptome analysis at 2 h and 6 h following mitochondrial morphology change induction. After switching on the iCMM system in iCMM^*mYF*^, iCMM^*YF*^, iCMM^*Cav1s*^, and iCMM^*mActZ*^ cells, the expression levels of 25, 23, 65, and 49 genes at 2 h and of 6, 42, 58, and 111 genes at 6 h, respectively, were significantly altered (Supplementary Fig. 10a). Consistent with a previous report^49^, rapamycin treatment increased *CREB3* regulatory factor (*CREBRF*) in all cells expressing iCMM at 2 h, highlighting the fidelity of this analysis (Supplementary Table 1). For 13 genes, mRNA levels were changed in the same direction and with similar timing by all functional iCMM effectors (Supplementary Table 1). Conversely, mRNA level changes were specific for 18, 31, 68, and 111 genes in iCMM^*mYF*^, iCMM^*YF*^, iCMM^*Cav1s*^, and iCMM^*mActZ*^ cells, respectively (Supplementary table 1).

When compared with iCMM^*mYF*^ cells, 0, 26, and 53 genes at 2 h and 39, 66, and 92 genes at 6 h were significantly differentially expressed after switching on the iCMM system in iCMM^*YF*^, iCMM^*Cav1s*^, and iCMM^*mActZ*^ cells, respectively (Supplementary Fig. 10b). mRNA levels of six genes were changed in the same direction in all cells expressing a functional iCMM effector, whereas the expression of 20, 58, and 98 genes was specifically altered in iCMM^*YF*^, iCMM^*Cav1s*^, and iCMM^*mActZ*^ cells, respectively (Supplementary Table 2). Collectively, these results indicated that the transcriptome was, at least partially, coordinated with mitochondrial morphology.

Gene set enrichment analysis of 50 “hallmark” gene sets representing major biological processes^50,51^ revealed that certain gene sets, including MYC targets and oxidative phosphorylation genes, were significantly enriched upon mitochondrial morphology change in cell lines expressing a functional iCMM effector (Fig. 6a). Compared with the reference iCMM effector mYF, each functional iCMM effector exhibited a distinctive hallmark pattern (Supplementary Fig. 11a). Gene ontology (GO) analysis revealed that iCMM-induced mitochondrial morphology change led to the reorganization of various cellular functions, most notably, ribosomal functions, in a time- and functional iCMM effector-dependent manner (Fig. 6b, Supplementary Fig. 11b). Regulatory target analysis showed that certain target genes, including *ELK1, NRF2*, and *SRF* were enriched, especially in cells expressing mActZ-FY (Fig. 6c, Supplementary Fig. 11c). These results indicated that mitochondrial morphology changes were reflected in the transcriptome within a few hours following their induction, and that they evoked various cellular functions.

**Fig. 6:**
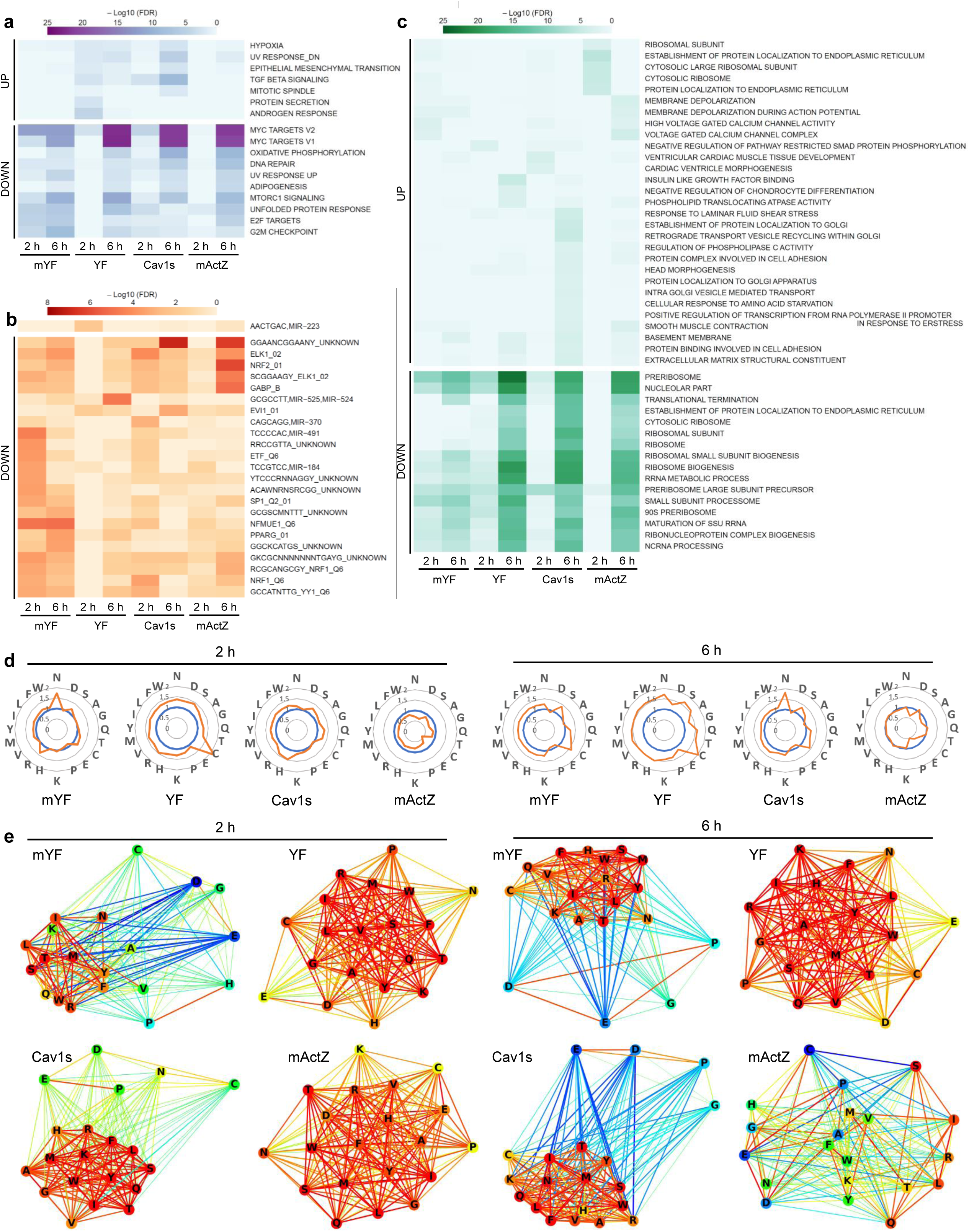
iCMM-induced mitochondrial morphology change results in reorganization of the transcriptome and amino acid profiles in cells. **a–c**, Hallmark gene sets (a), GO terms (b), and regulatory targets (c) enriched upon mitochondrial morphology change in cells stably expressing iCMM were analyzed (n = 3), and heatmaps of – Log10 (FDR) are shown. Cells were harvested at the indicated time points after adding DMSO (iCMM: OFF) or rapamycin (iCMM: ON). Darker color indicates lower FDR. The FDR was calculated from comparisons between DMSO- and rapamycin-treated cells for each effector. Up/Down indicates the direction of a transcription change in rapamycin-treated compared to DMSO-treated cells. Up to a maximum of 30 gene sets with FDR < 0.01 in a direction and category are shown. **d, e**, Amino acid profiles of cells stably expressing iCMM were visualized (n = 3). Cells were harvested at the indicated time points after adding DMSO (iCMM: OFF) or rapamycin (iCMM: ON). (d) Radar charts of the amino acid profiles of cells stably expressing iCMM. Blue: DMSO-treated cells, orange: rapamycin-treated cells. (e) Network graphs based on the correlation coefficient for each amino acid. The color of each node represents the correlation with each amino acid based on serine. The color of the edge represents the correlation between amino acids. Blue: low correlation coefficient, red: high correlation. The thickness of the edge represents the absolute value of the correlation coefficient.

### Rapid mitochondrial morphology change by iCMM alters the amino acid profile

Of the 20 amino acids utilized for protein synthesis, 17 require mitochondrial enzymes for their metabolism^52^. We examined whether mitochondrial morphology change affected the intracellular amino acid profile in HeLa cells stably expressing iCMM. In iCMM^*mYF*^ and iCMM^*Cav1s*^ cells, each amino acid displayed different changes upon iCMM system activation (Fig. 6d). In contrast, nearly all amino acids tended to increase upon mitochondrial morphology change in iCMM^*YF*^ cells, whereas in iCMM^*mActZ*^ cells, they tended to decrease (Fig. 6d). Findings were similar at 2 h and 6 h after mitochondrial morphology change induction (Fig. 6d). To analyze the interplay between mitochondrial morphology and amino acid profile further, we examined correlations among amino acids. Network architectures reflecting amino-acid correlations were similar between iCMM^*mYF*^ and iCMM^*Cav1s*^ cells, and between iCMM^*YF*^ and iCMM^*mActZ*^ cells (Fig. 6e). Glutamate, aspartate, and proline, which are regulated in p53-mediated amino acid metabolism in the vicinity of mitochondria^53^, presented a low correlation with serine, which is critical for mitochondrial dynamics and functions^54^, in iCMM^*mYF*^ and iCMM^*Cav1s*^ cells (Fig. 6e). However, in iCMM^*YF*^ cells, nearly all amino acids presented a strong correlation with serine, whereas correlations among amino acids in iCMM^*mActZ*^ cells disappeared over time (Fig. 6e). These results suggested that the amino acid profile is altered in accordance with the mitochondrial morphology change induced by iCMM.

## Discussion

While accumulating evidence highlights the importance of mitochondrial morphology in various cellular functions, the mechanisms by which it orchestrates a dynamic intracellular signaling network in harmony with various subcellular compartments remain poorly understood. We developed iCMM as a powerful tool to manipulate mitochondrial morphology in a living cell on a minutes timescale. As the intracellular mitochondrial morphology distribution is determined by the demand of mitochondria at the subcellular compartment level^55^, iCMM provides an advantage in resolving the biological significance of mitochondrial morphology, which would otherwise have been challenging. We established three iCMM systems capable of inducing different mitochondrial morphologies, and we investigated the biological significance of each mitochondrial morphology using these systems.

We found that, even after mitochondrial morphology change, the ΔΨm remained stable for several hours, suggesting that there exists a system that protects the ΔΨm in the unlikely event that mitochondrial morphology is altered by single or multiple environmental signals. However, mitochondrial morphology change triggered various biological responses, including changes in the OCR, mitochondrial parkin recruitment, and intracellular omics information. This suggests that mitochondrial morphology changes are not random, but are purposely coordinated based on varied cellular information. If this is true, in future, it may be possible to extract intracellular information profiles from mitochondrial morphology information using a computational approach.

The advantage of iCMM—similar to other CID system-based devices—is that mitochondrial morphology can be altered in different ways by changing the effector and/or anchor. mitochondrial morphology change can be induced based on another principle by using a different effector, whereas the anchor can be designed so that the effector functions only in a specific part of the mitochondria (defined by the anchor localization), allowing fine-tuning of the mitochondrial morphology change even when the same effector is used. As mitochondria exhibit different morphologies, depending on the physiological situation, it is important to create more iCMM systems for comprehensive investigation of the biological significance of mitochondrial morphology.

Decoding biological information embedded in mitochondrial morphology is one of the fascinating avenues in the field of mitochondrial biology. By demonstrating the usefulness of iCMM, we corroborated the biological significance of mitochondrial morphology in various cellular functions, suggesting the possibility of constructing a system that can purposefully manipulate cellular functions by changing the mitochondrial morphology. Indeed, transcriptome and amino acid profiles were reorganized upon iCMM-mediated mitochondrial morphology manipulation. Further studies will hopefully unravel the physiological significance of mitochondrial morphology in various cellular functions.

In summary, the iCMM system developed in this study allows effective and rapid manipulation of mitochondrial morphology. When combined with conventional genetic approaches, iCMM may provide new insights into the physiopathological functions of mitochondrial morphology in health and disease. Moreover, the working principle could be applied to other organelles. The system holds promise for improving our understanding of the biological significance of cellular organelle morphology. Consolidating precise manipulation of organelle morphology with clinical medicine will provide a novel therapeutic approach to optimizing drug efficacy.

## Methods

### Plasmid construction

The mYF effector was produced via point mutation of EYFP, by replacing Ala^206^ with Lys in the YF vector (Addgene, #20175), using the Q5 Site-Directed Mutagenesis Kit (New England Biolabs, E0554S). The YF-Cav1s effector was produced by subcloning the sequence coding aa 61–178 of human caveolin 1 (UniprotKB-Q03135) into a YF vector at the C-terminus between *Hind*III and *Sal*I. The mActZ-FY effector was produced by subcloning sequences coding aa 30–262 of codon-optimized *Listeria monocytogenes* serovar 1/2a ActA (UniprotKB-P33379) and aa 2–380 of human zyxin (UniprotKB-Q15942) into an FY vector containing the FKBP^F100Y^ mutant and EYFP, at the N-terminus between the *Xho*I and *EcoR*I sites. The ActA nucleic acid sequence was optimized for mammalian cells^24^. The Tom20-CR anchor was produced by inserting a stop codon encoding Tom20-CR^56^ after the *Xho*I site in the vector, using the Q5 Site-Directed Mutagenesis Kit. ER and Golgi apparatus-specific marker proteins were generated by subcloning sequences coding aa 100–134 of rat cytochrome b5 (UniprotKB-P00173) and aa 3131–3259 of human golgin subfamily B member 1 (UniprotKB-Q14789) into the mCherry vector at the C-terminus between the *EcoR*I and *Sal*I sites, respectively. To generate lysosome-specific marker proteins, the sequence coding aa 1–417 of human LAMP1 (UniprotKB-P11279) harboring V119A and L170P mutations was subcloned into the mCherry vector at the N-terminus, between the *Nhe*I and *Age*I sites. To obtain CRISPR-Cas9 expression targeting the *AAVS1* locus, donor vectors containing Tom20-CR driven by the *CAG* promoter were constructed using pAAVS1-P-CAG-mCh (Addgene, #80492). The pAAVS1-P-CAG-mCh vector was inversely amplified using primers that excluded the mCherry sequence flanked by the *EcoR*I site (termed pAAVS1-P-CAG-Tom20-CR). The insert coding Tom20-CR was amplified, followed by the assembly of the vector and insert using In-Fusion cloning (TaKaRa). For lentiviral expression of effector constructs, each effector was amplified and then subcloned into the *EcoR*I and *Sal*I sites of the pLenti-EF-IRES-blast vector (a gift from Yutaka Hata). All constructs were verified by sequencing following subcloning.

### Cell culture and transfection

Human cervical adenocarcinoma HeLa cells (CCL-2), human hepatocellular carcinoma Hep 3B cells (HB-8064), and human osteosarcoma U2-OS cells (HTB-96) were purchased from the American Type Culture Collection and were cultured in Dulbecco’s modified Eagle’s medium (DMEM; Thermo Fisher, 11965118) supplemented with 10% fetal bovine serum (FBS; Thermo Fisher, 10270-106) and 1% Zell Shield (Minerva Biolabs GmbH, 13-0050) at 37 °C in 5% CO_2_. Torin 1 was purchased from Merck (475991). For transient transfection of the iCMM system, 2.4 × 10^5^ cells were plated on a poly-lysine-coated glass-bottom dish (Matsunami, D1131H) and incubated for 2 h at 37 °C in 5% CO_2_. Following incubation, the cells were transfected with the plasmid using FuGENE HD (Promega, E2311). Indicated experiments were carried out 20–32 h after transfection.

### Lentivirus production

HEK293T cells were transiently transfected with pLenti-EF-blast vectors together with pCAG-HIVgp and pCMV-VSV-G-RSV-Rev (provided by RIKEN BRC, Ibaraki, Japan) using TransIT-2020 Reagent (TaKaRa). The medium, containing lentivirus, was collected.

### Establishment of HeLa cells stably expressing the iCMM system

First, a HeLa cell line that stably expressed Tom20-CR was established. For gene targeting at the *AAVS1* locus, pXAT2 and pAAVS1-P-CAG-Tom20-CR were transfected into 1 × 10^6^ cells in a single-cell suspension using NEPA21 electroporation. Two days after electroporation, 2 μg ml^−1^ of puromycin was added with daily feeding over a period of 7 days to select for targeted cells. Serial dilution cloning was performed to isolate single-cell-derived clones. The clone was incubated with lentivirus encoding an iCMM effector in the presence of 5 μg ml^−1^ polybrene. After 48 h, the clones were cultured in the presence of 5 μg ml^−1^ blasticidin for 10 days to select for cells expressing the iCMM effector. Following the selection process, cloning was performed using 3.2-mm sterile cloning discs (Merck).

### Live-cell imaging

ECFP, EYFP, and mCherry excitation was carried out using an Intensilight mercury-fiber illuminator (Nikon). Data were processed through a CFP-A-Basic-NTE filter (Semrock), YFP-A-Basic-NTE filter (Semrock), and mCherry-B-NTE-ZERO filter (Semrock) for ECFP, EYFP, and mCherry imaging, respectively. Cells were viewed using a 40× objective (Plan Apochromat Lambda Series, Nikon) mounted on an inverted Eclipse Ti2-E microscope (Nikon) and imaged using a Zyla 4.2 PLUS sCMOS camera (Oxford Instruments). Imaging data were processed using the NIS-Elements AR 5.01 imaging software (Nikon). All imaging experiments were completed at 37 °C in 5% CO_2_ using an STX stage top incubator (Tokai-Hit). For all live-cell imaging, cells were suspended in phenol red-free DMEM (Thermo Fisher, 31053028) supplemented with 10% FBS, 4 mM L-glutamine (Thermo Fisher, 25030081), and 1% penicillin-streptomycin (Sigma-Aldrich, P4333). Time was measured from the first frame (0 min), and 50 nM rapamycin (Calbiochem) was added at the indicated time.

### mitochondrial morphology analysis

Cells were cultured for 30 min in the presence of 0.5 μM MitoTracker Red CM-H2Xros (Thermo Fisher, M7513) at 37 °C in 5% CO_2_. The cells were washed with imaging medium twice, and the fluorescence in mitochondria was recorded using NIS software (Nikon). Fluorescence images were processed using Unsharp Mask (Power: 1.0; Area: 41) followed by Rolling Ball Correction (Radius 1.95 µm; bright signal) using NIS-Elements AR 5.01. Subsequently, the image was processed by contrast limited adaptive histogram equalization, followed by thresholding using the open-source software Fiji (22743772). Average mitochondrial footprint, average number of individual mitochondria, and roundness of mitochondria were analyzed using the ‘Analyze Particle’ function in Fiji.

### CLEM

CLEM was carried out at Japan Electron Optics Laboratory (Tokyo, Japan). Briefly, HeLa cells (cultured on a glass-bottom dish) that transiently expressed the iCMM system were washed twice with 1 ml of PBS. The cells were then fixed for 10 min in a fixation mixture containing 4% formaldehyde (NEM, 3153) and 0.1% glutaraldehyde (NEM, 304) in PBS at room temperature. After three washes with PBS, the cells were imaged using a Nikon A1R equipped with a Nikon A1plus camera, an Apo 40× WI λS DIC N2 (Nikon), as well as a 405 and a 408 laser for ECFP and EYFP excitation, respectively. For scanning electron microscopy (SEM) imaging, cells were subjected to post-fixation (1% OsO_4_ and 1% tannic acid), Bloc contrast staining (1% uranyl acetate and lead aspartate), and dehydration, followed by Epon embedding. SEM imaging was carried out using a JSM-7900F (JEOL).

### ΔΨm analysis

Cells were cultured for 15 min in the presence of 50 nM TMRE at 37 °C in 5% CO_2_. They were then washed with imaging medium twice, and TMRE fluorescence in mitochondria was recorded with the NIS software (Nikon). Background fluorescence was measured in a cell area devoid of mitochondria and subtracted from the fluorescence obtained from mitochondria.

### Extracellular flux analysis

The OCR was measured using a Seahorse XF24 extracellular flux analyzer (Seahorse Bioscience). HeLa cells stably expressing the iCMM system were seeded in a 24-well culture plate (Seahorse Bioscience) at 5 × 10^4^ cells per well in 250 μl of culture medium and were incubated at 37 °C and 5% CO_2_ for 24 h. The cells were treated with DMSO or 50 nM rapamycin for the indicated time, and the culture medium was replaced with 525 µl of XF Base Medium pH 7.4 (Seahorse Bioscience) supplemented with 4 mM GlutaMAX and 25 mM D-glucose. The cells were incubated at 37 °C in a non-CO_2_ incubator for 30 min. Meanwhile, an XF24 sensor cartridge (hydrated overnight; Seahorse Bioscience) was loaded with the appropriate volumes of oligomycin (final concentration, 1 μM), protonophore FCCP (final concentration, 0.125 μM), and rotenone/antimycin A (final concentrations, 0.5 μM). Three basal oxygen consumption measurements were recorded (each for 8 min) before the addition of oligomycin, FCCP, and finally, rotenone/antimycin A. The effects of these chemicals on mitochondrial oxygen consumption were also measured three times, each for 8 min. Data were normalized to the protein concentration in each group, which was determined using the Protein Assay BCA Kit (Nacalai, Japan)

### Cell proliferation assay

Cells (1 × 10^6^) were cultured for 48 h at 37 °C in 5% CO_2_. Trypan blue (Thermo Fisher, 15250-061) stained cells were counted using a LUNA cell counter (Logos Biosystems).

### Transcriptome analysis

Cells stably expressing indicated iCMM system were harvested at 2 h or 6 h after rapamycin treatment, and RNA was isolated using Direct-zol 96 (Zymo Research). RNA-seq libraries were prepared from 500 ng of RNA using Quant-seq 3’ FWD (Lexogen) following the manufacturer’s instructions. An equal amount of each Quant-seq library was pooled and diluted to 4 pM. The library was denatured, and 2.3 nM of denatured library was subjected to RNA-seq in an Illumina Next-seq 500 instrument using high-output flow cells and the 75 single-end mode. Using BBDuk, low-quality bases were trimmed, and poly-A or -T sequences, adapter sequences, as well as 11-base and one-base reads (from the left and right sides of reads, respectively), were mapped to the human genome (GRChg38.p12) using STAR-aligner^57^. Read counts were obtained using featureCounts^58^. Samples that generated at least one million mapped reads were used for further analyses. Downstream analysis was performed using R [R Core Team (2017). R: A language and environment for statistical computing. R Foundation for Statistical Computing, Vienna, Austria. URL https://www.R-project.org/]. Differentially expressed genes (DEGs) were identified using edgeR^59,60^. To determine DEGs, we employed a generalized linear model, and the gene-wise likelihood ratio test. DEGs with fold change > 2 and FDR-adjusted P < 0.05 were considered significant. Competitive gene set tests were performed to account for inter-gene correlation using the camera function in the limma package as well as three gene-set collections (hallmark, C3 regulatory target, and C5 GO) from the Molecular Signatures Database (MSigDB)^50,51,61^. An FDR < 0.01 was considered significant, and up to 30 gene sets in each direction (up- or downregulated) of a gene-set collection were reported. If both directions occurred in two or more comparisons, the FDR was set as 1 for the different direction (e.g., GO_Actomyosin was enriched in upregulated genes in mActZ, but it also enriched in downregulated genes in other comparisons. In the plot showing upregulated genes, FDRs for the comparisons other than mActZ are 1).

### Metabolome analysis

Cells growing on 10-cm dishes were washed twice with ice-cold PBS and intracellular metabolites were extracted by briefly incubating the cells with 1 ml of methanol containing internal control substances (50 μM 2-morpholinoethanesulfonic acid and 50 μM methionine sulfone) on ice. Cell debris was removed by centrifugation (14,000 × *g* for 10 min at 4 °C), and 600 μl of supernatant was mixed with 300 μl of ultrapure water and 450 μl of chloroform. Following centrifugation (16,000 × *g* for 3 min at 4 °C), 800 μl of supernatant was mixed with 400 μl of ultrapure water and centrifuged again. The supernatant (1 ml) was evaporated for 40 min to reduce the organic solvent content. The samples were subjected to ultrafiltration using 3-kDa filters (Amicon Ultra 3K device, Merck). After lyophilization, the samples were dissolved in 50 μl of ultrapure water. Metabolome analysis was conducted by LC-MS/MS (LCMS-8030, Shimadzu, Kyoto, Japan) using the primary metabolite method package version 2 (Shimadzu) according to the manufacturer’s protocol.

### Apoptosis analysis

Cells were treated with DMSO or rapamycin for 1 h and then incubated with 266 nM STS for 6 h. Then, the cells were incubated with Hoechst 33342 (Thermo Fisher) for 10 min for nuclear staining. After fixation with a 4% paraformaldehyde phosphate buffer solution (Wako, 163-20145), the cells were scored as possessing normal or apoptotic nuclei, in several fields. Three independent experiments were conducted. Data are reported as the percentage of cells with apoptotic nuclei among total cells.

### Western blot analysis

Total cell lysates were prepared using NP40 cell lysis buffer (Thermo Fisher, FNN0021) containing a protease/phosphatase inhibitor cocktail (Cell Signaling Technology, #5872), and protein concentrations were determined using a BCA assay (Thermo Fisher). For western blotting, protein samples were separated by sodium dodecyl sulfate-polyacrylamide gel electrophoresis and transferred to nitrocellulose membranes (Bio-Rad, 1620115). The membranes were blocked in Tris-buffered saline-Tween 20 containing 5% nonfat milk at room temperature for 1 h. The membranes were then incubated with the following primary antibodies for 18 h at 4 °C: anti-caspase-3 (#14220), anti-phospho-p70 S6 kinase (Thr389) (#9234), anti-p70 S6 kinase (#2708), and anti-β-actin (#4970) (Cell Signaling Technology). The membranes were subsequently incubated with horseradish peroxidase-conjugated goat anti-rabbit IgG (Cell Signaling Technology) and visualized using chemiluminescence detection (Immobilon, Millipore). Coomassie brilliant blue staining was conducted using Bullet CBB Stain One Super (Nacalai, 13542-65) according to manufacturer’s protocol.

### Amino acid correlation network analysis

The network graph was drawn using the Fruchterman–Reingold algorithm. This uses a force-directed graph-drawing algorithm to determine the position of each node from the relationship between the nodes and the edges. The nodes receive the force and moves. A force that brings the node closer (attractive force) and a force that moves the node away (repulsive force) work simultaneously. The attractive force *f*_*a*_ and the repulsive force *f*_*r*_ were defined by the following equations:

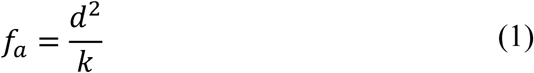

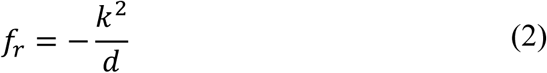

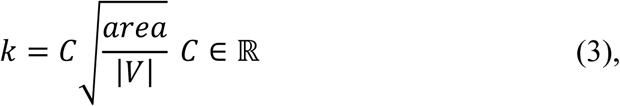

where d was the distance between nodes, area was the area of the drawing space, and | V | was the number of nodes. The force applied to the node was *f*_*a*_ + *f*_*r*_, and is shown in the graph below (k = 10).

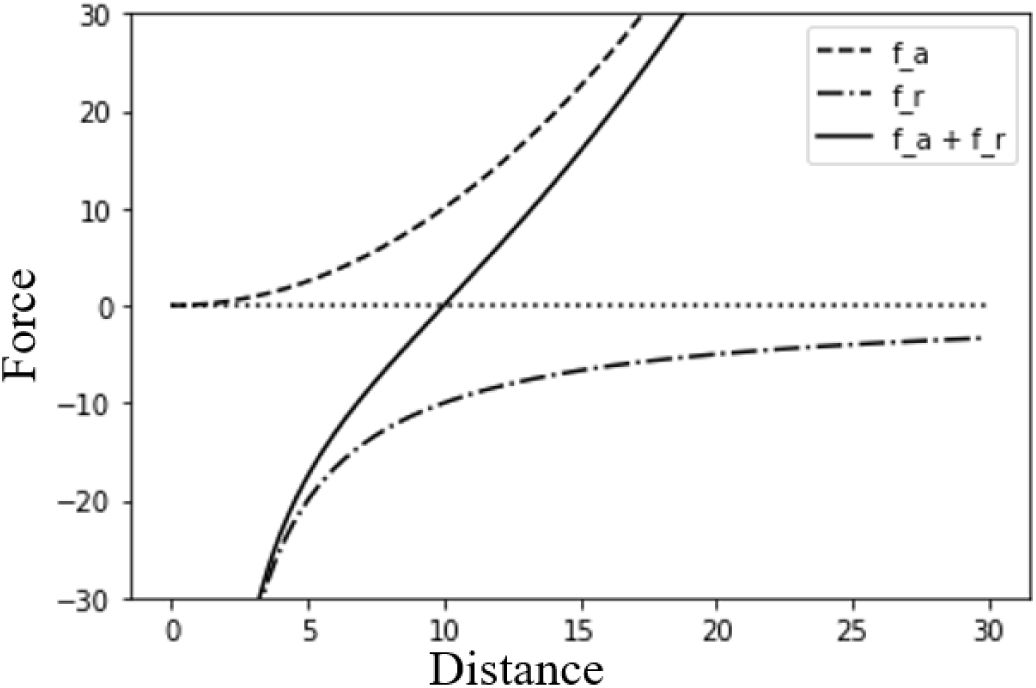

In addition, the temperature *t* that limits the amount of movement of the node was lowered step by step. The moving direction vector is given by Equation 4.

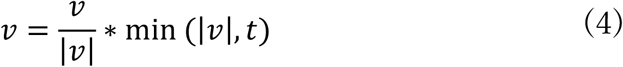

The initial position of each node was randomly assigned, the positions of the nodes were updated using the force and temperature parameters, and this calculation was repeated a certain number of times. The development environment was python 3.7, and the graph was created using NetworkX version 2.3. The edge weights were given by the correlation coefficient of each amino acid. The k value of the graph-drawing parameters of NetworkX was set to 1.5, and the other parameters used default values.

### Statistical analysis

Statistical analysis was performed using an paired t-test and unpaired two-tailed Student’s t test. The F test was used to determine whether variances were equal or unequal.

## Acknowledgements

We thank Katsuyuki Suzuki, Akira Takebe (JEOL Ltd) and Katsuko Okubo (University of Tsukuba) for technical assistance, Keita Takahashi (NikonInstech CO., Ltd), Hiroyuki Soejima (Tokyo Science CO., Ltd), and Yasuko Maruyama (University of Tsukuba) for generous support. This study was supported in part by Grant-in-Aid for Scientific Research on Innovative Areas (18H04854), Leading Initiative for Excellent Young Researchers, JST CREST (JPMJCR1927).

## Author contributions

T.Miyamoto. conceived the project. T.Miyamoto. designed the experiments. T.Miyamoto., H.U., Y.M., S.G., D.Y., M.M., Y.Y., H.N., N.H., Y.T., M.S., T.Matsuzaka, F.H., SI.T., N.Y., K.I., H.S. conducted the experiments. T.Miyamoto. and H.U. wrote the manuscript.

---

**Supplementary Figure 1.**
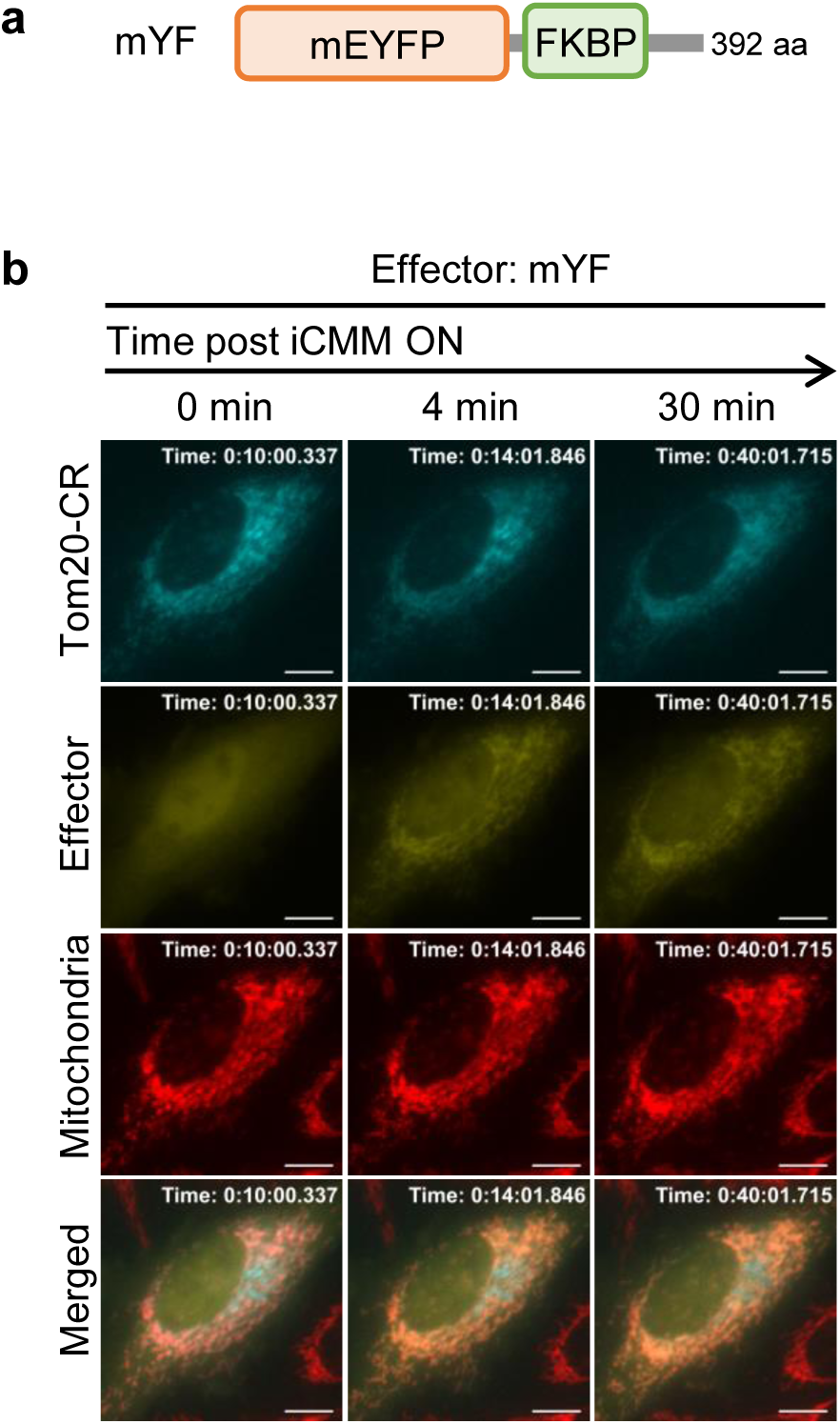

**Supplementary Figure 2.**
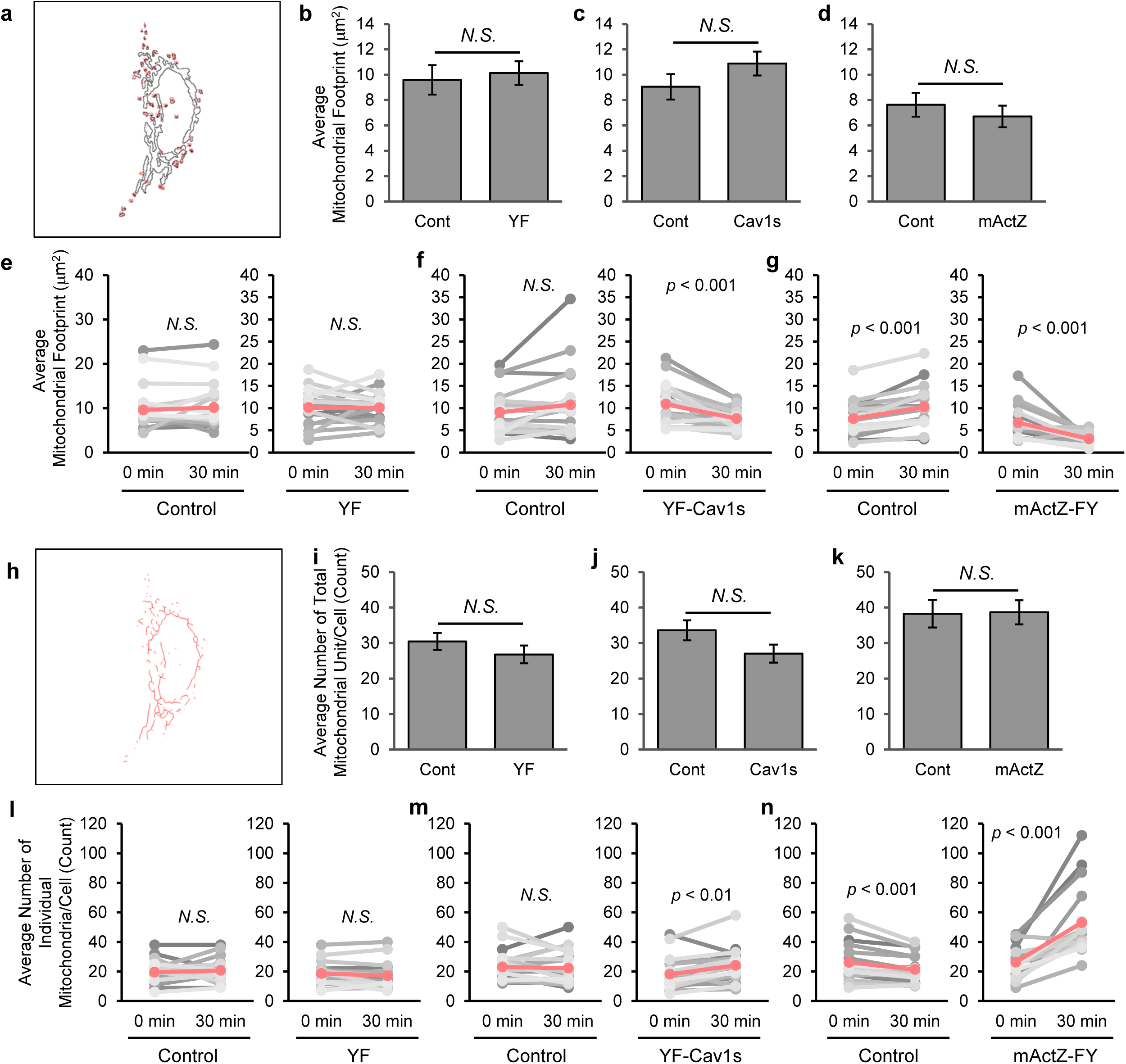

**Supplementary Figure 3.**
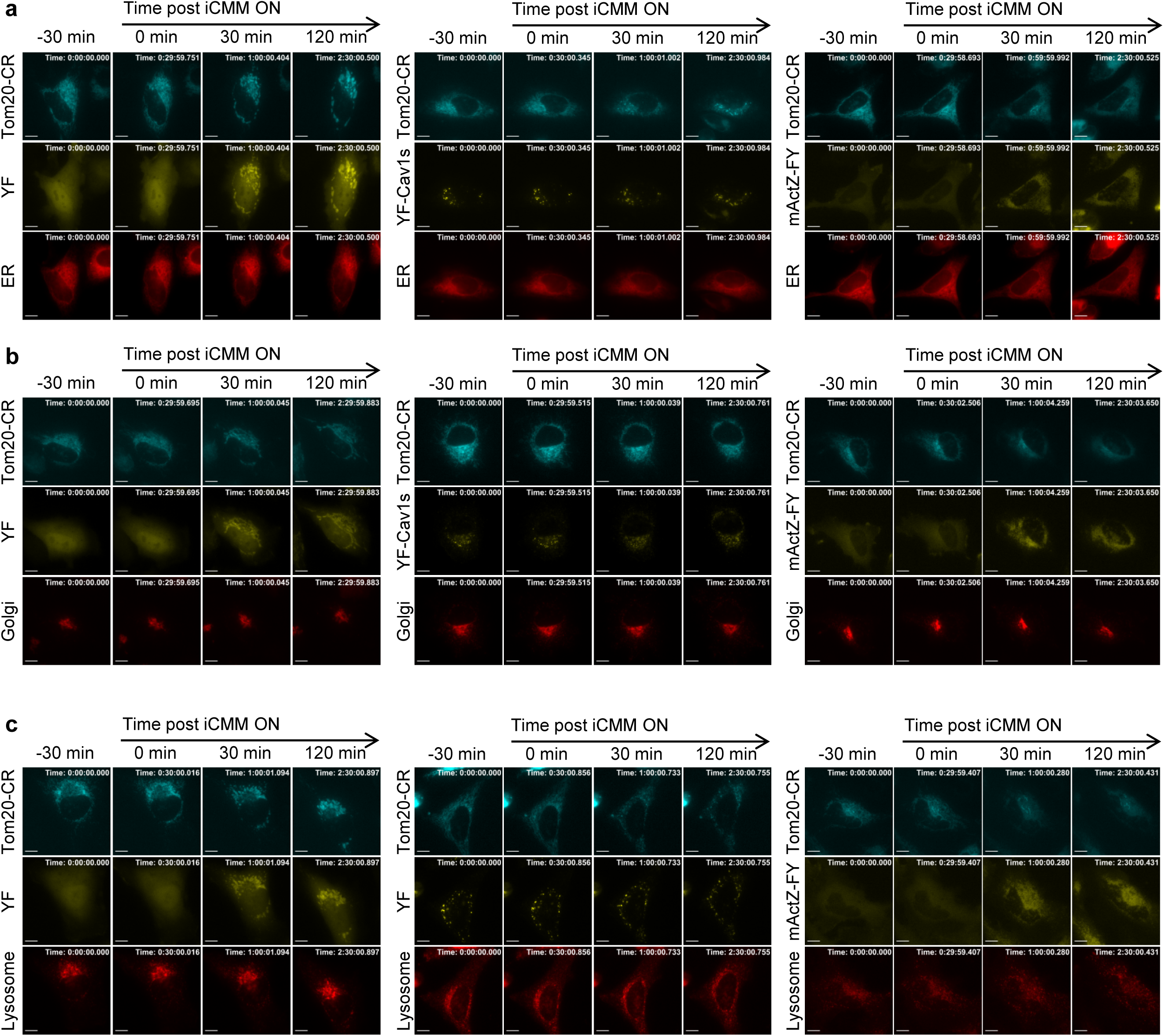

**Supplementary Figure 4.**
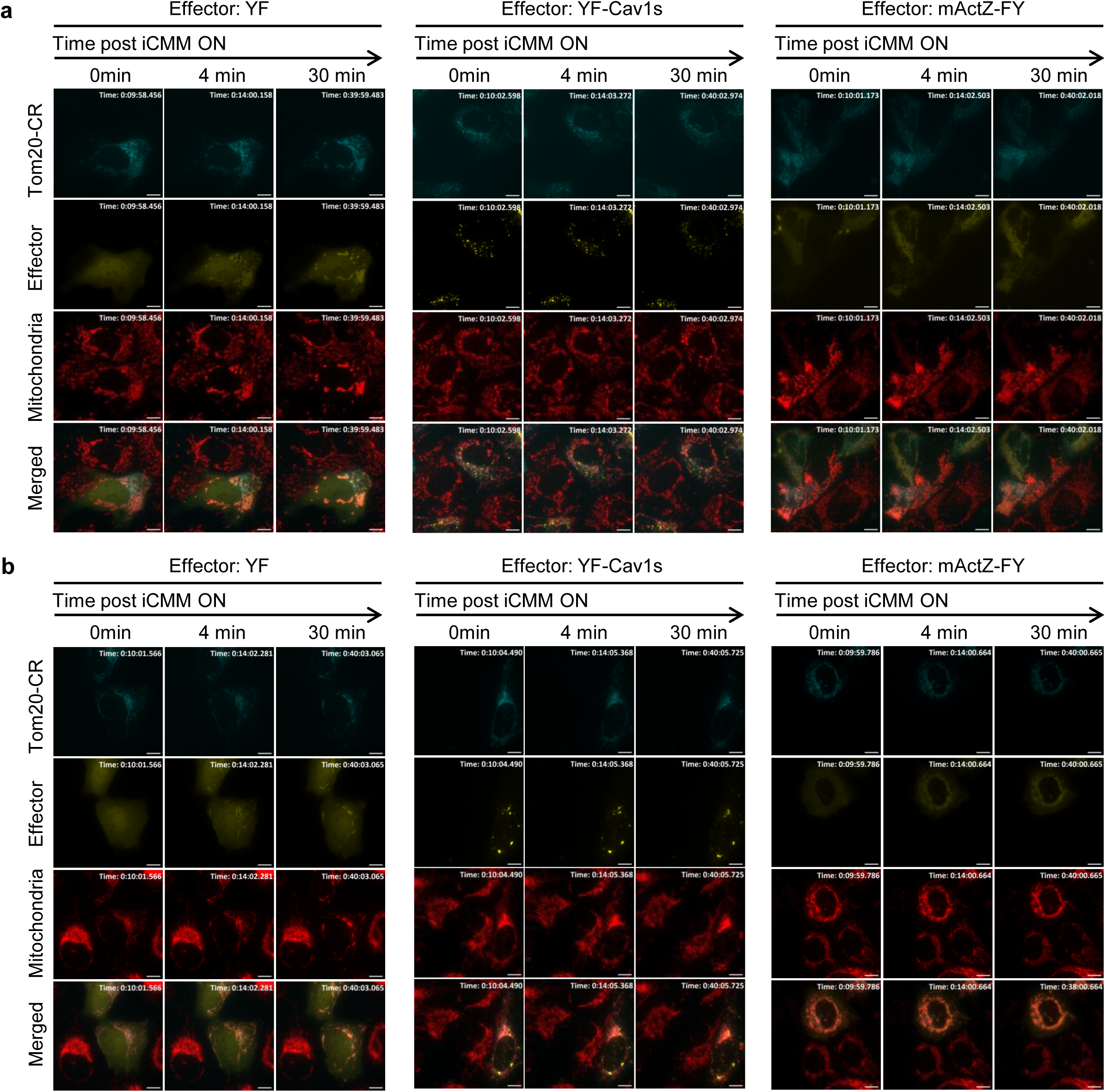

**Supplementary Figure 5.**
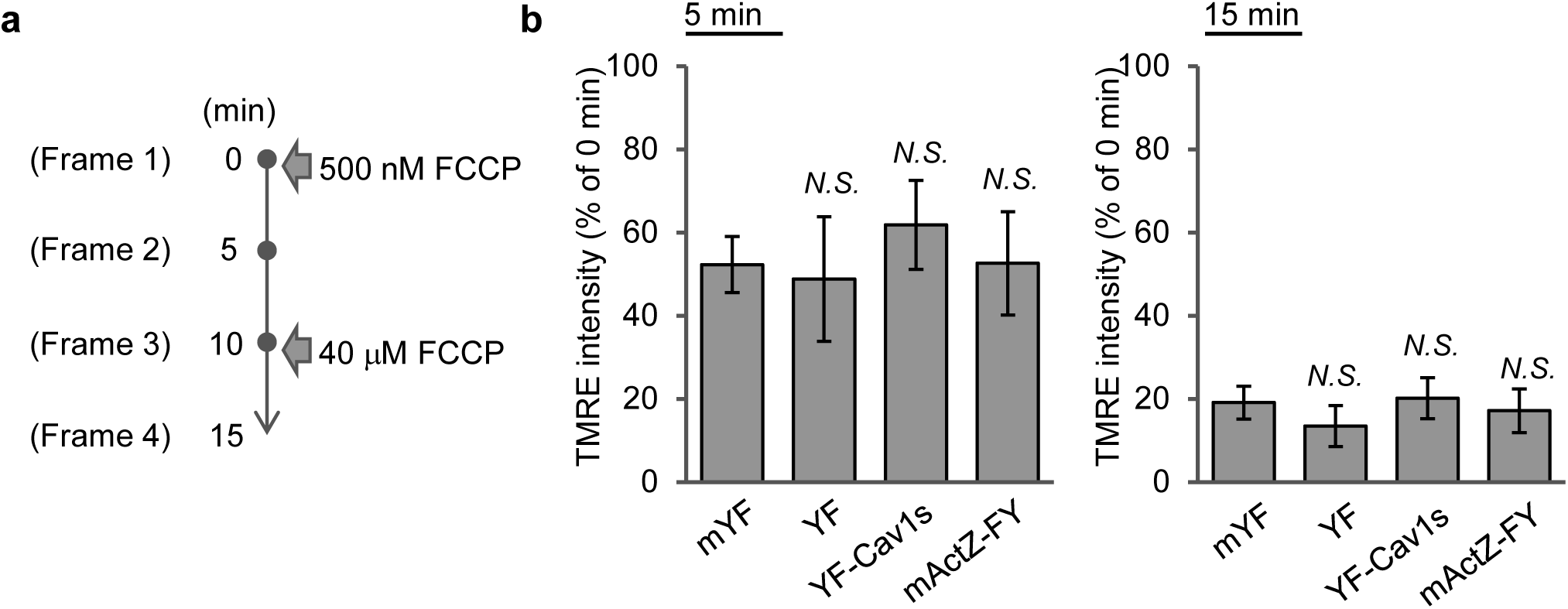

**Supplementary Figure 6.**
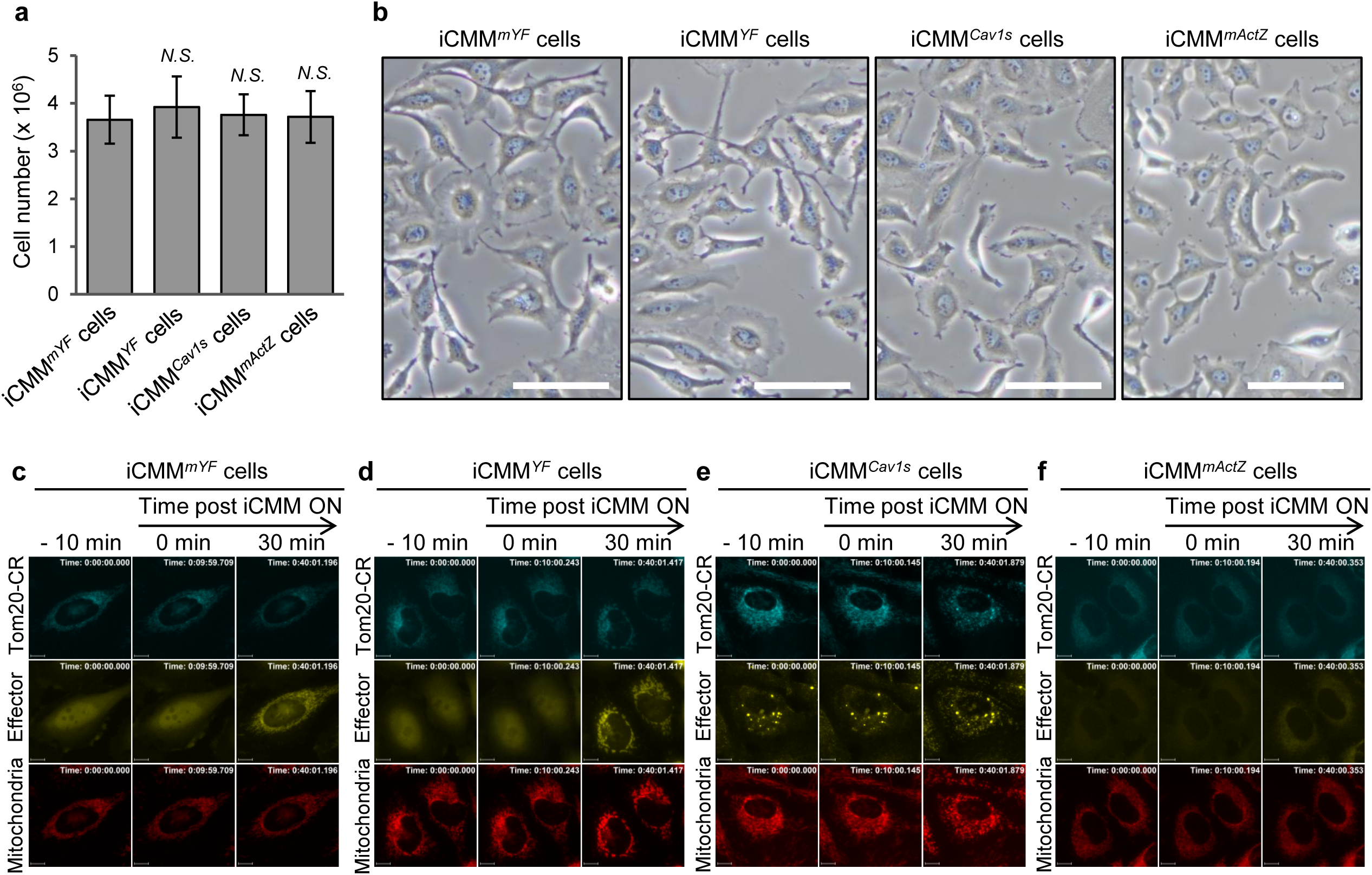

**Supplementary Figure 7.**
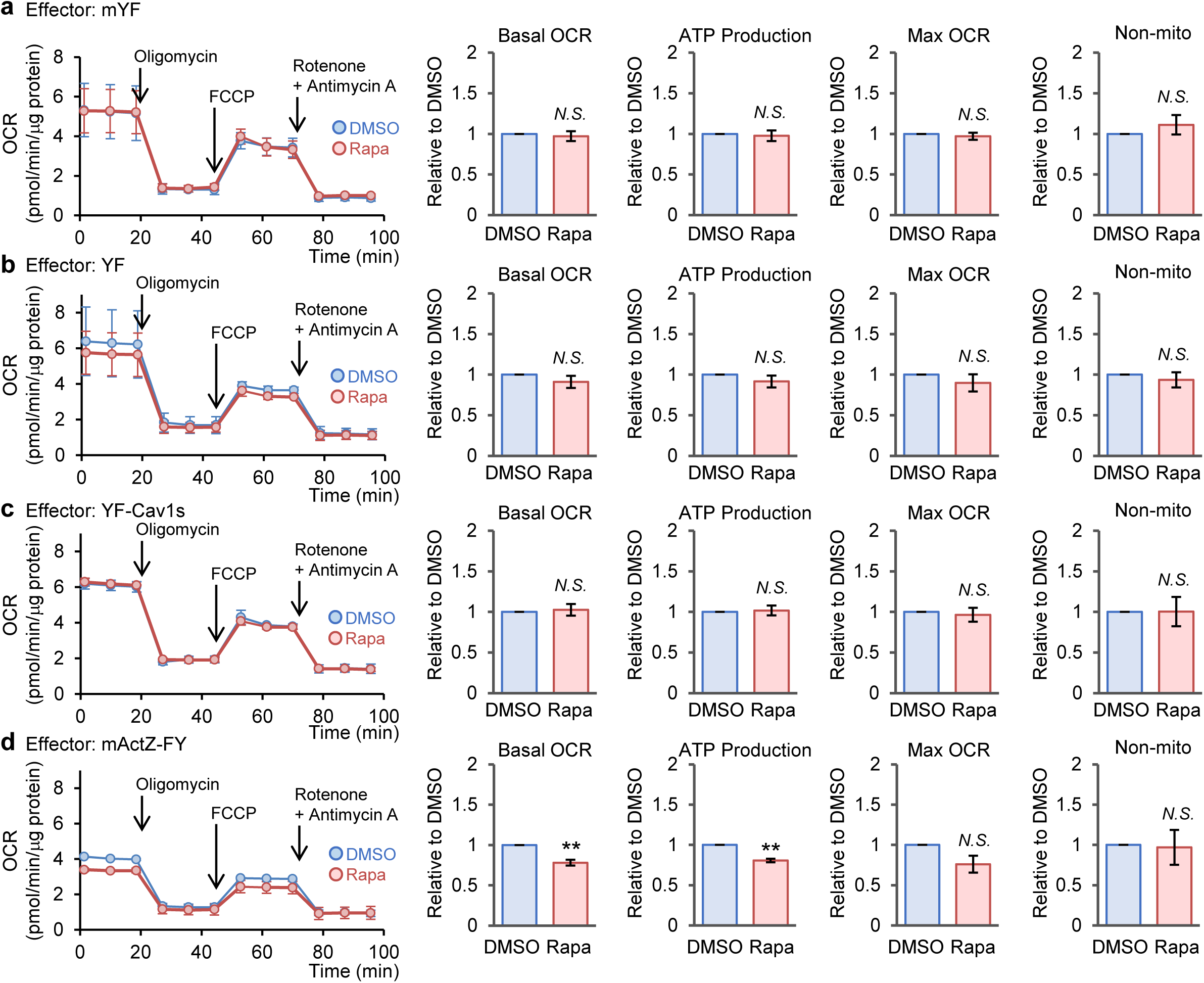

**Supplementary Figure 8.**
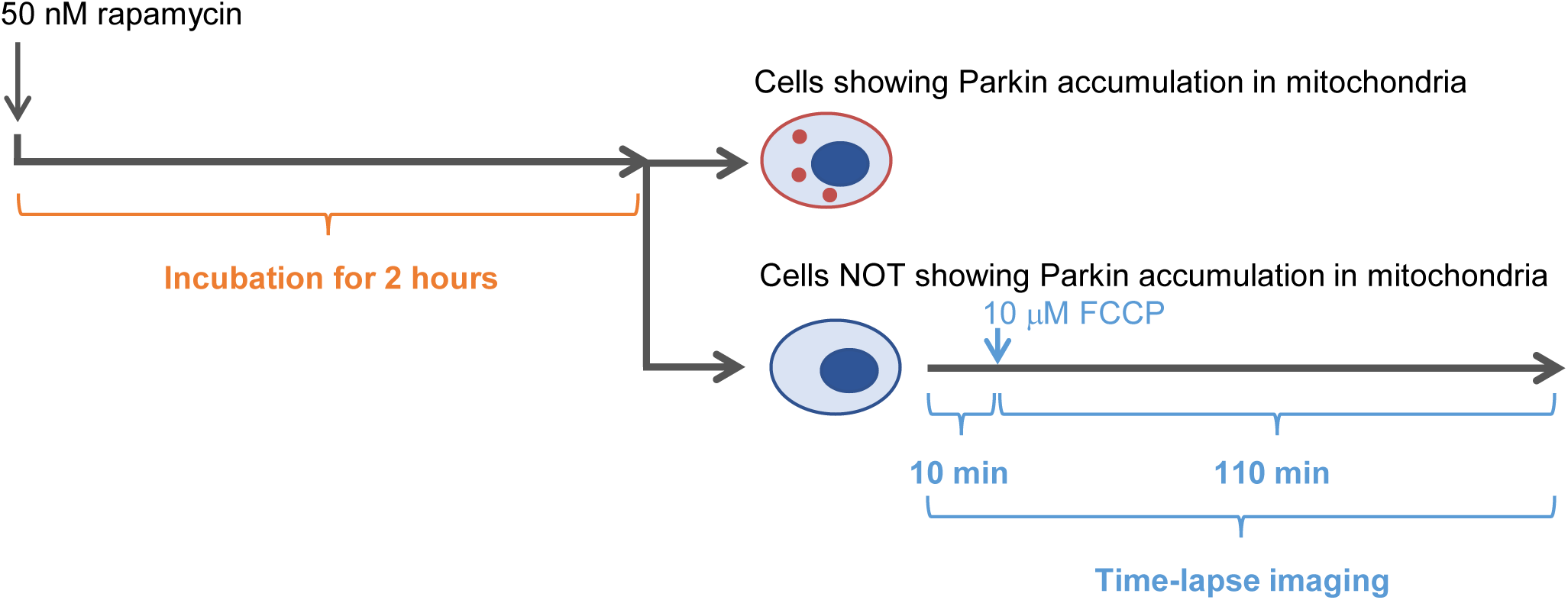

**Supplementary Figure 9.**
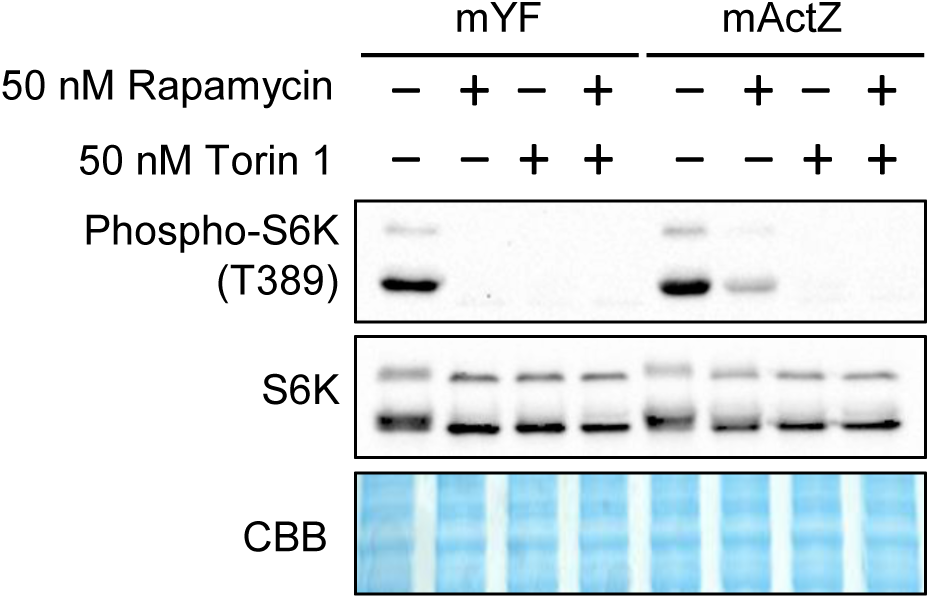

**Supplementary Figure 10.**
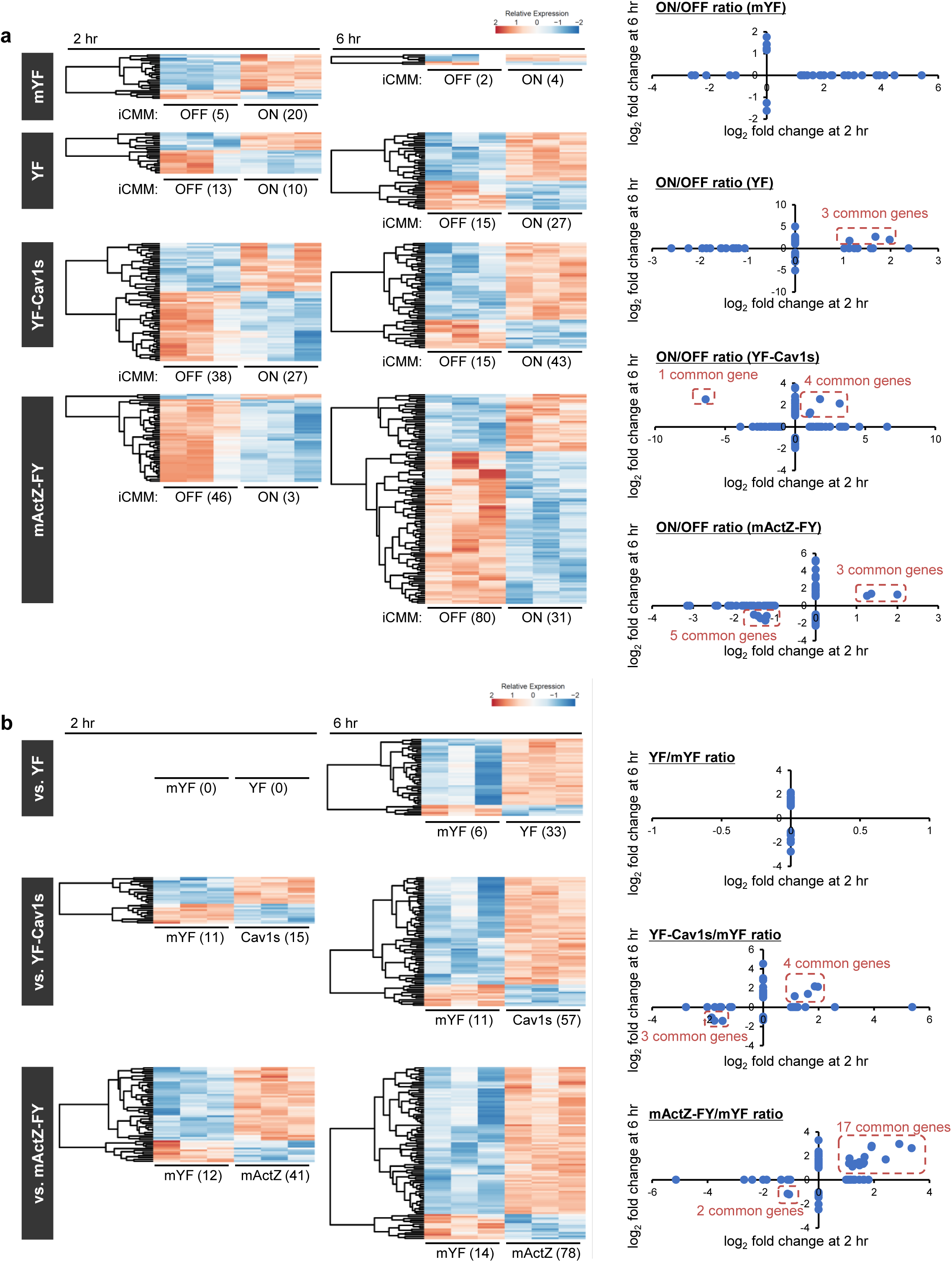

**Supplementary Figure 11.**
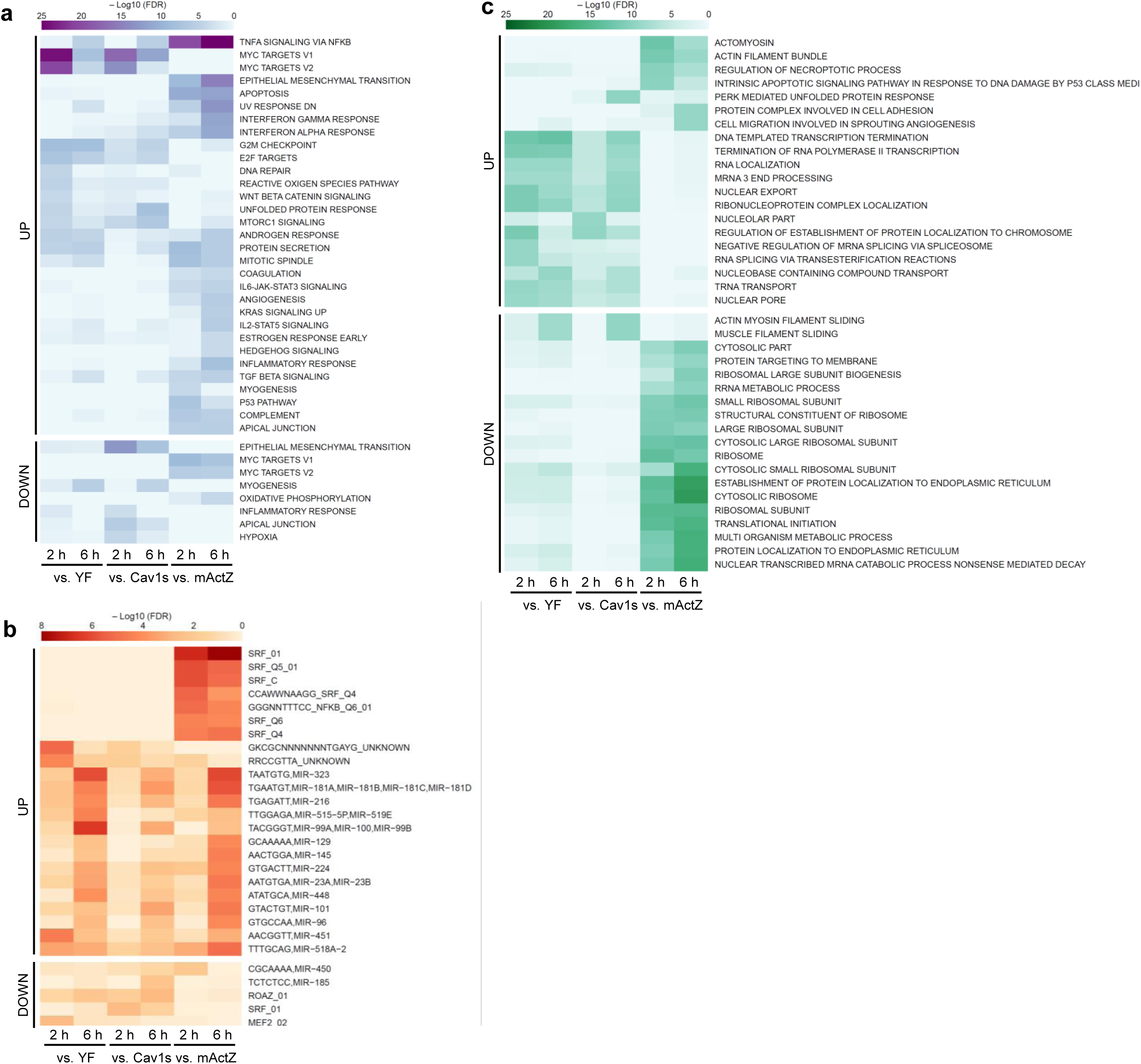

