## Supplementary information for "Disentangling the biological information encoded in disordered mitochondrial morphology through its rapid elicitation by iCMM"

**Supplementary Fig. 1: mYF as a control effector for iCMM.**

**a**, Structural features of mYF. **b**, Time-lapse images of HeLa cells expressing Tom20-CR and mYF. Mitochondria (red) were stained with MitoTracker Red CMXRos. Scale bar: 10 μm.

**Supplementary Fig. 2: Effect of iCMM on mitochondrial area and number.**

**a**, Representative images of mitochondria for mitochondrial area analysis. **b–d**, Mitochondrial areas in HeLa cells expressing the indicated iCMM system and in surrounding iCMM-negative cells (control) were analyzed before adding rapamycin. Data are presented as the mean ± standard deviation. Cell data sets in Supplementary Fig. 2b, 2c, and 2d are similar to those in Fig. 1f, 1g, and 1h, respectively. *N.S.*: statistically nonsignificant (Student's t-test). **e–g**, Mitochondrial areas in HeLa cells expressing the indicated iCMM system and in surrounding iCMM-negative cells (control) were analyzed before (0 min) and after (30 min) rapamycin treatment. Gray: analyzed single cell, red: average. $p$ = paired t-test. *N.S.*: statistically nonsignificant. Cell data sets in Supplementary Fig. 2e, 2f, and 2g are similar to those in Fig. 1f, 1g, and 1h, respectively. **h**, Representative images of mitochondria for mitochondrial number analysis. **i–k**, Mitochondrial numbers in HeLa cells expressing the indicated iCMM system and in surrounding iCMM-negative cells (control) were determined before adding rapamycin. Data are presented as the mean ± standard deviation. Cell data sets in Supplementary Fig. 2i, 2j, and 2k are similar to those in Fig. 1f, 1g, and 1h, respectively. *N.S.*: statistically nonsignificant (Student's t-test). **l–n**, Mitochondrial numbers in HeLa cells expressing the indicated iCMM system and in surrounding iCMM-negative cells (control) were determined before (0 min) and after (30 min) rapamycin treatment. Gray: analyzed single cell, red: average. $p$ = paired t-test. *N.S.*: statistically nonsignificant. Cell data sets in Supplementary Fig. 2l, 2m, and 2n are similar to those in Fig. 1f, 1g, and 1h, respectively.

**Supplementary Fig. 3: Effect of iCMM on non-target organelle morphology.**

**a–c**, Representative time-lapse images of HeLa cells expressing the indicated iCMM system and organelle marker. ER: mCh-Cb5 (a), Golgi: mCh-GOLGB1 (b), lysosome: mCh-LAMP1 (c). Scale bar: 10 μm.

**Supplementary Fig. 4: Versality of iCMM.**

**a, b**, Time-lapse images of Hep3B (a) and U-2 OS (b) cells expressing iCMM. Mitochondria (red) were stained using MitoTracker Red CMXRos. Scale bar: 10 μm.

**Supplementary Fig. 5: Interplay between changes in mitochondrial morphology and sensitivity to FCCP.**

**a**, Time course of the experiment performed in Fig. 2e–h. **b**, Quantitative data for Fig. 2e–h. All data are presented as the mean ± standard deviation. *N.S.*: statistically nonsignificant (Student's t-test).

**Supplementary Fig. 6: Establishment of HeLa cells stably expressing iCMM.**

(a) Indicated $1 \times 10^6$ cells were cultured for 48 h, and the total cell number was determined. Quantification was performed in three independent experiments. All data are presented as the mean ± standard deviation. *N.S.*: statistically nonsignificant (vs. iCMM$^{mYF}$ cells, Student's t-test). (b) Representative phase difference images of the indicated cells. (c) Time-lapse images of HeLa cells stably expressing iCMM systems. Mitochondria (red) were stained using MitoTracker Red CMXRos; 2 min/frame. Rapamycin was added after image acquisition at frame 6 (indicated as "0 min"). Scale bar: 100 μm.

**Supplementary Fig. 7: Effect of prolonged mitochondrial morphology change by iCMM on the OCR.**

**a–d**, OCR in iCMM$^{mYF}$ (a), iCMM$^{YF}$ (b), iCMM$^{Cav1s}$ (c), and iCMM$^{mActZ}$ (d). Before OCR measurement, each cell line was treated with DMSO or rapamycin for 120 min. All data are presented as the mean ± standard deviation from three independent experiments. *$P < 0.05$, **$P < 0.01$; *N.S.*: statistically nonsignificant (paired t-test).

**Supplementary Fig. 8: Schematic diagram of the Parkin translocation experiment.**

The experimental procedure for Fig. 4 is shown.

**Supplementary Fig. 9: Effects of rapamycin and torin 1 on mTORC1 activity.**

HeLa cells stably expressing Tom20-CR and the indicated iCMM effector were treated with rapamycin or torin 1 for 6 h. The cells were then subjected to western blot analysis.

**Supplementary Fig. 10: Heatmap of differentially expressed genes (DEGs) in HeLa cells stably expressing iCMM.**

**a**, Heatmaps and graphs of DEGs between DMSO and rapamycin treatments at 2 h and 6 h. (b) Heatmaps and graphs of DEGs between mYF and other effectors with rapamycin treatment at 2 h and 6 h. It should be noted that there were no DEGs between mYF and YF at 2 h. The normalized expression of each gene is shown in red (higher expression) and blue (lower expression). Genes were clustered according to Euclidean distances, and dendrograms represent the distances. The number of DEGs is shown in parentheses.

**Supplementary Fig. 11: Comparative transcriptome data analysis of the control iCMM effector**

**and a functional iCMM effector.**

**a–c**, Gene sets of hallmarks (a), gene ontology (b), and regulatory target (c) were analyzed and heatmaps of –Log10 (FDR) are depicted. Cells were harvested at the indicated time points after adding DMSO (iCMM: OFF) or rapamycin (iCMM: ON). Darker color indicates lower FDR. The FDR was calculated in comparisons between mYF and the indicated functional iCMM effector. Up/Down indicates the direction of a transcriptional change for the functional iCMM effector compared to mYF.

**Supplementary Table 1.**

List of differentially expressed genes before and after activating the iCMM system.

**Supplementary Table 2.**

List of differentially expressed genes after a change in mitochondrial morphology by a functional iCMM effector.

**Supplementary Movie 1.**

HeLa cells transiently expressing Tom20-CR and YF

**Supplementary Movie 2.**

HeLa cells transiently expressing Tom20-CR and mYF

**Supplementary Movie 3.**

HeLa cells transiently expressing Tom20-CR and YF-Cav1s

**Supplementary Movie 4.**

HeLa cells transiently expressing Tom20-CR and mActZ-FY
