## Supplementary figures for "Disentangling the biological information encoded in disordered mitochondrial morphology through its rapid elicitation by iCMM"

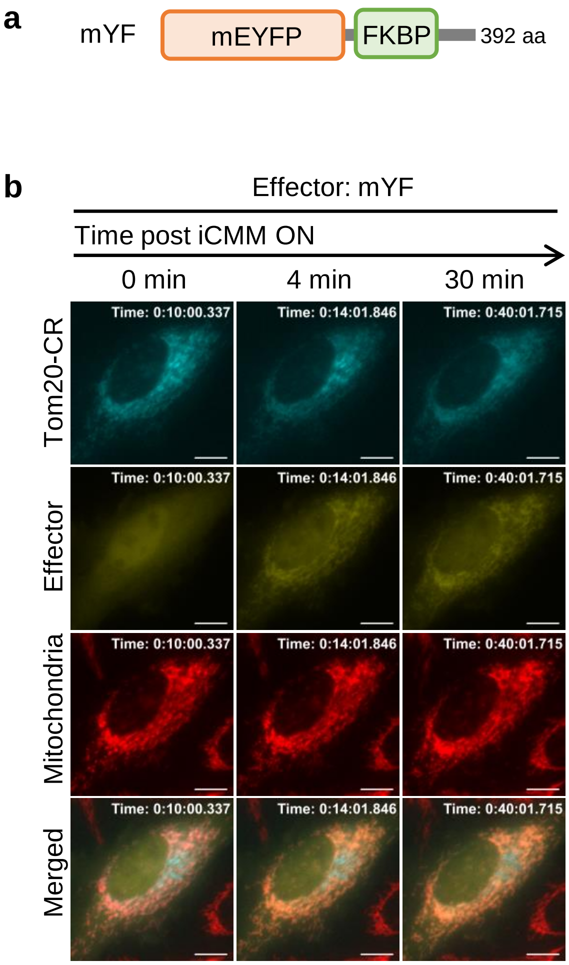


**Supplementary Figure 1. Miyamoto et al.**

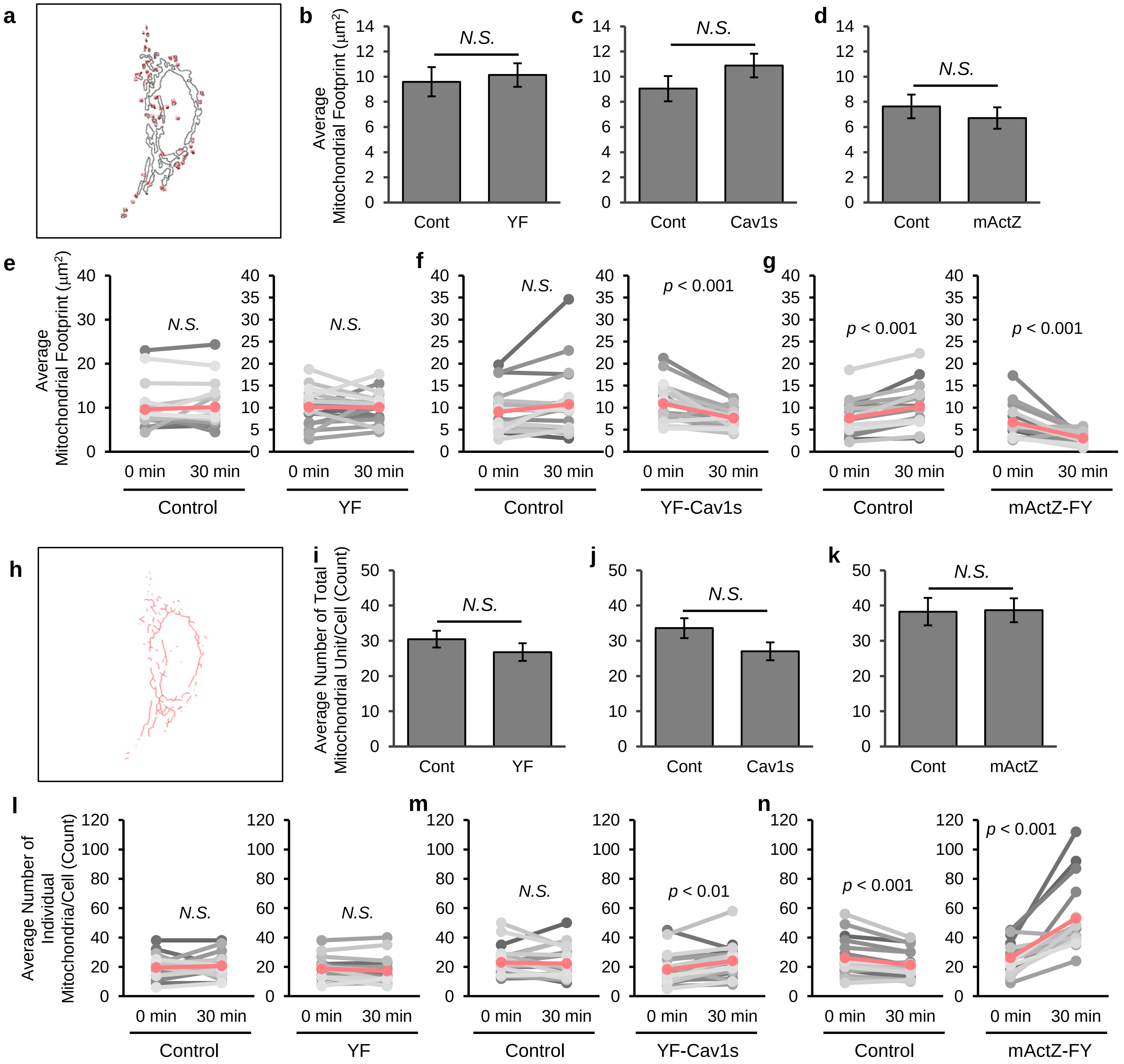


**Supplementary Figure 2. Miyamoto et al.**

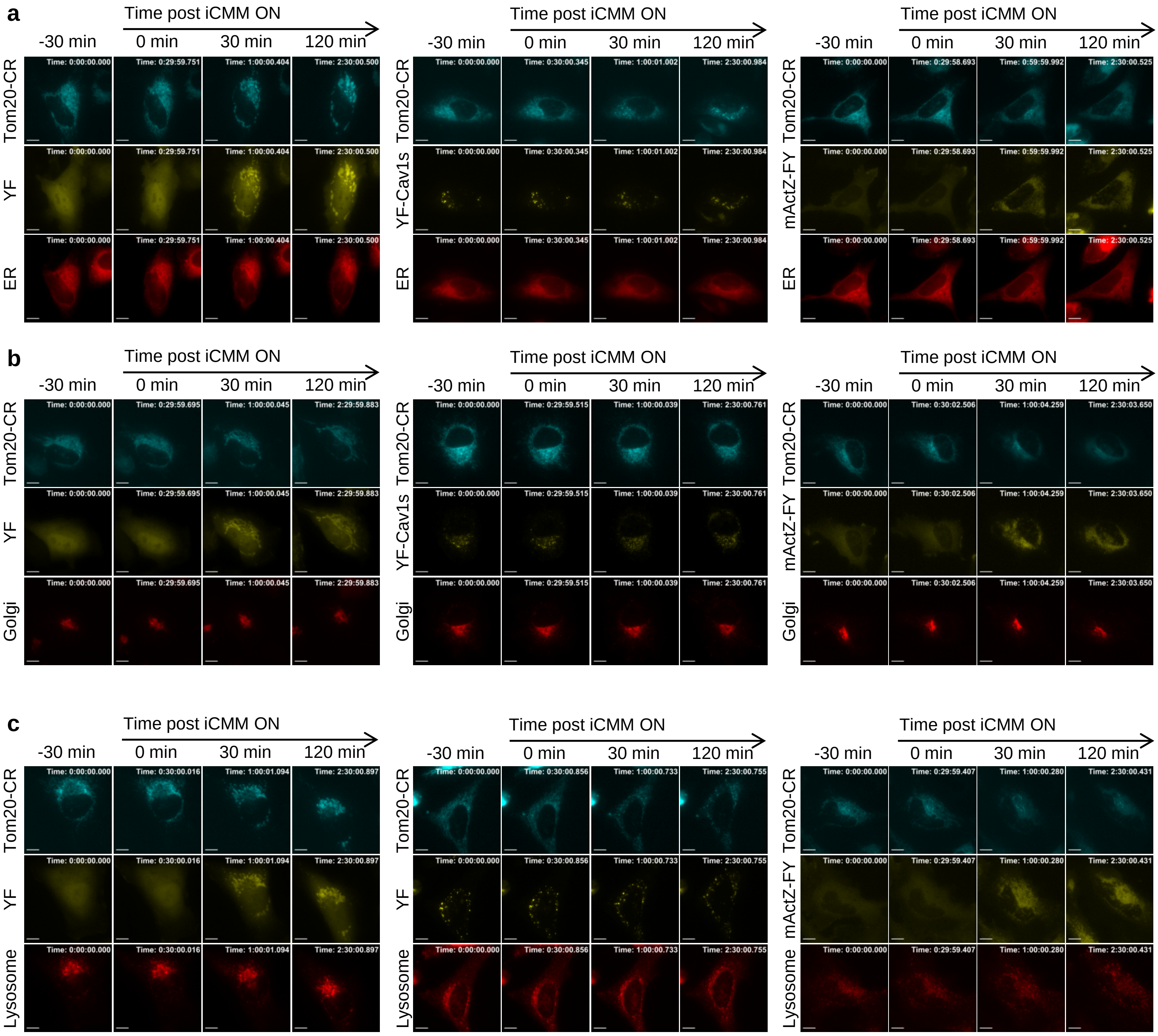


**Supplementary Figure 3. Miyamoto et al.**

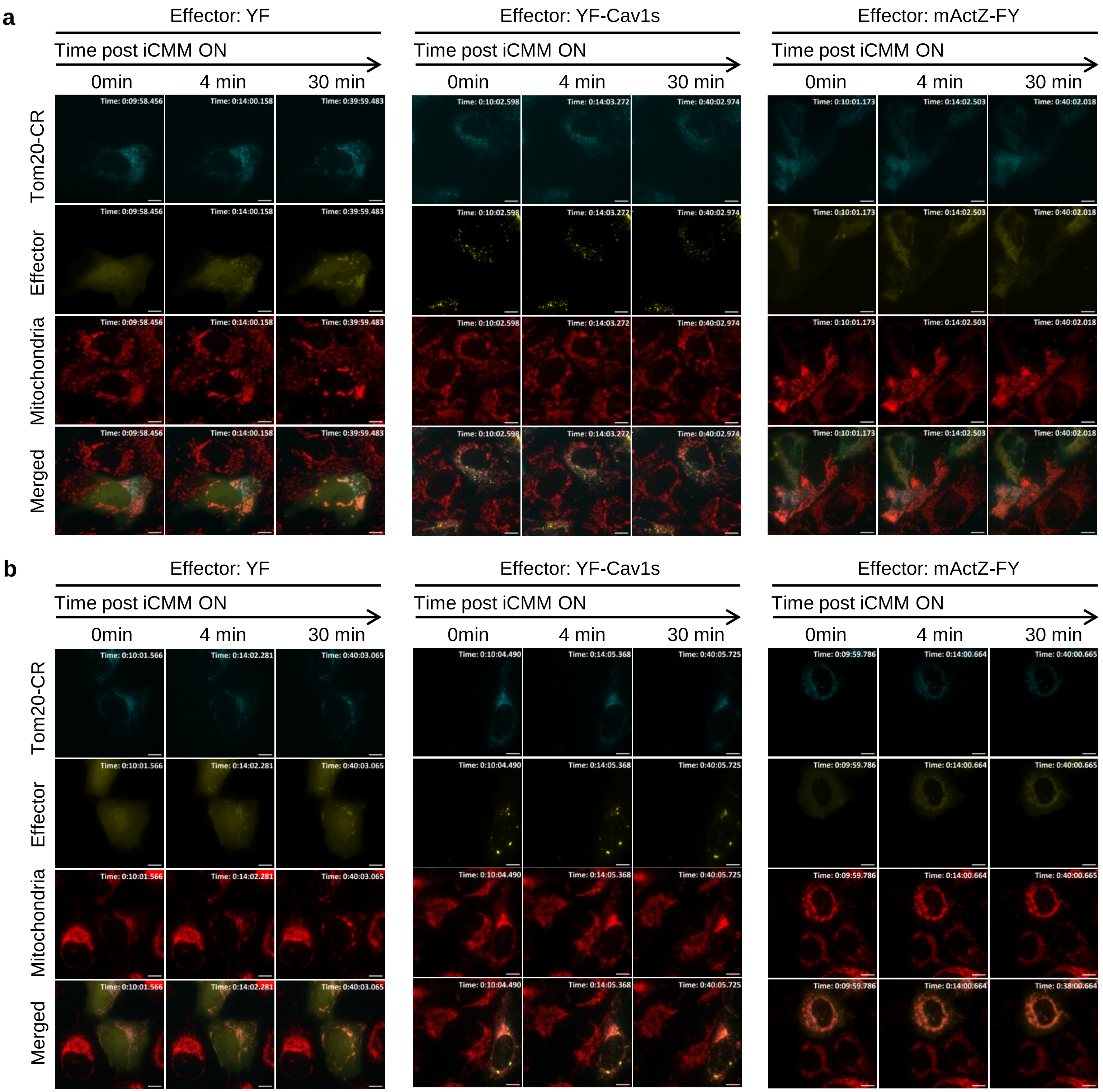


**Supplementary Figure 4. Miyamoto et al.**

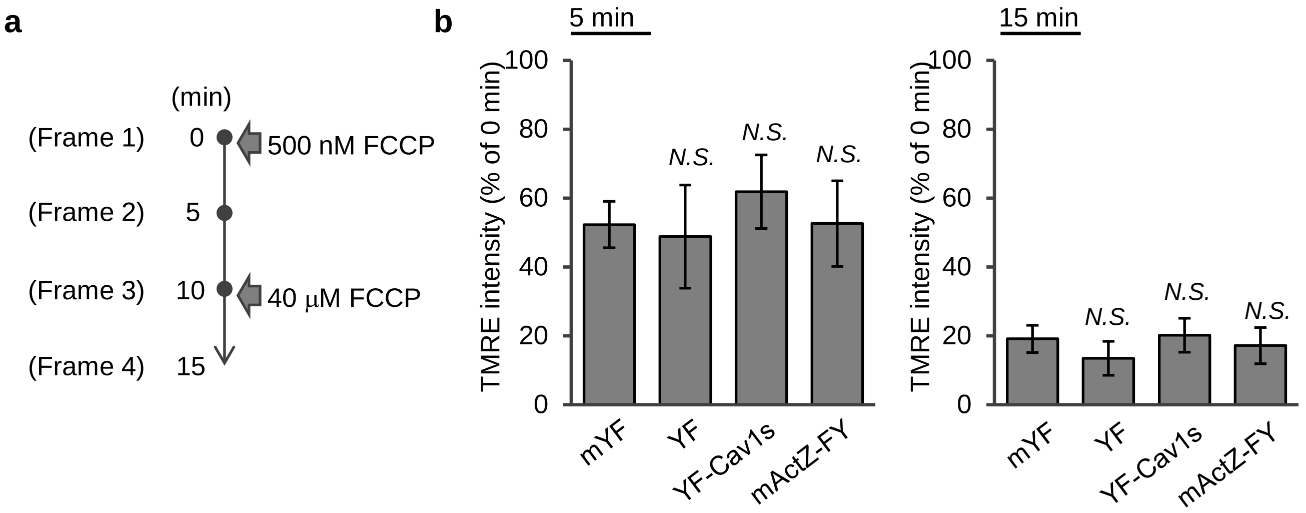


**Supplementary Figure 5. Miyamoto et al.**

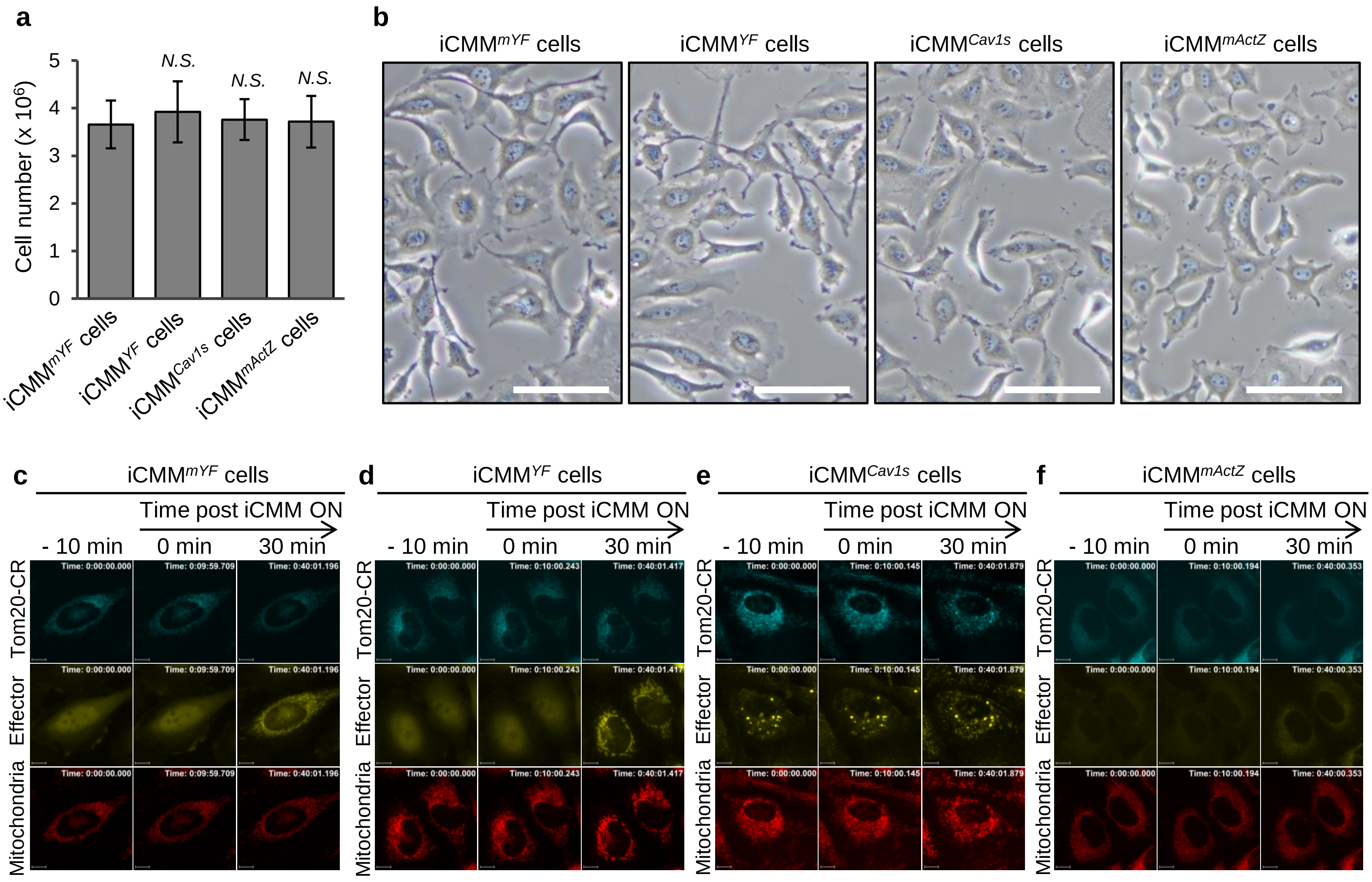


**Supplementary Figure 6. Miyamoto et al.**

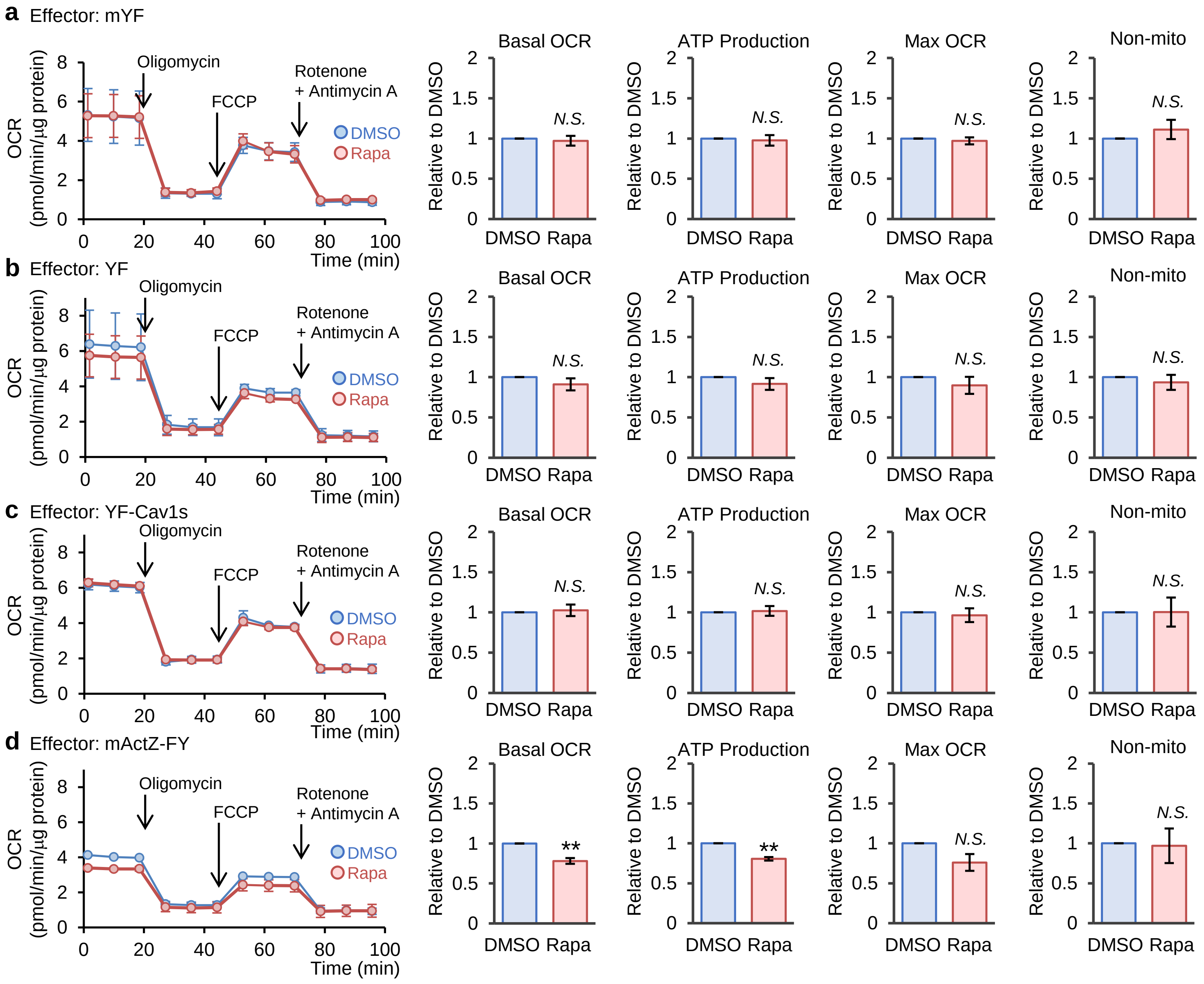
a Effector: mYF
b Effector: YF
c Effector: YF-Cav1s
d Effector: mActZ-FY
OCR
(pmol/min/μg protein)
Time (min)
Oligomycin
FCCP
Rotenone
+ Antimycin A
DMSO
Rapa
Basal OCR
ATP Production
Max OCR
Non-mito
Relative to DMSO
N.S.
**

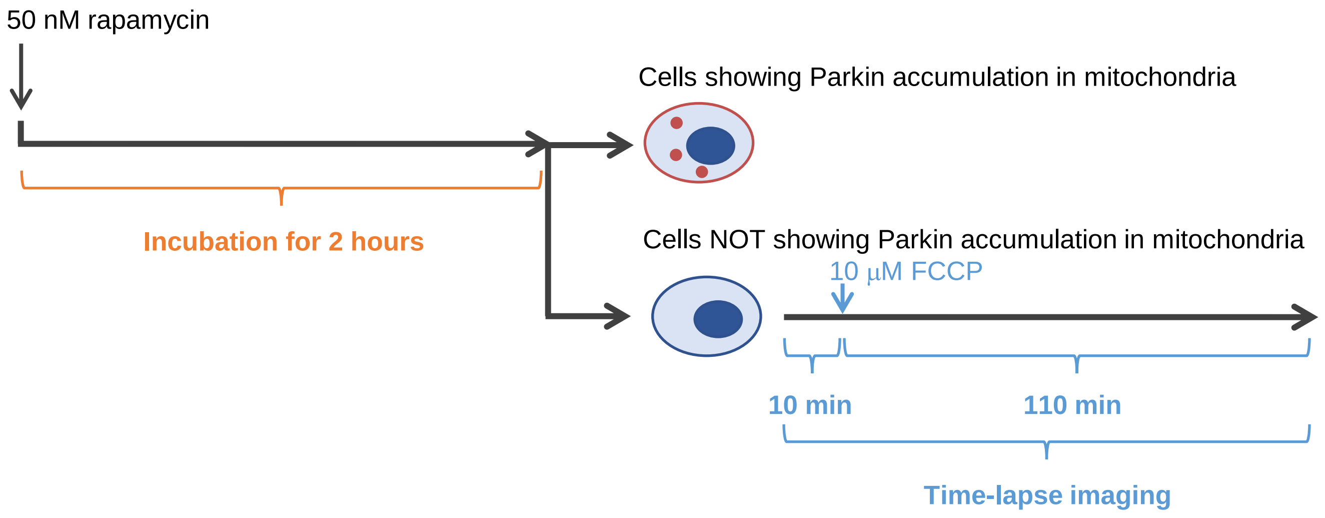


**Supplementary Figure 8. Miyamoto et al.**

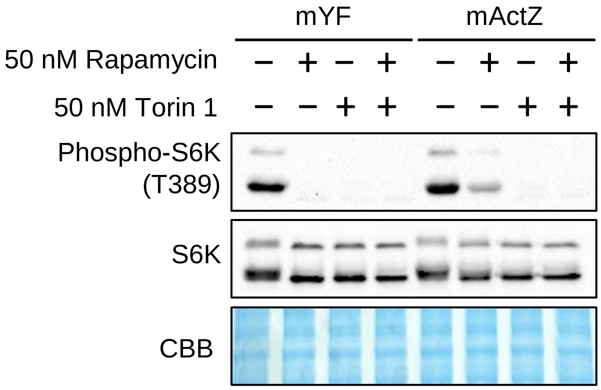


**Supplementary Figure 9. Miyamoto et al.**

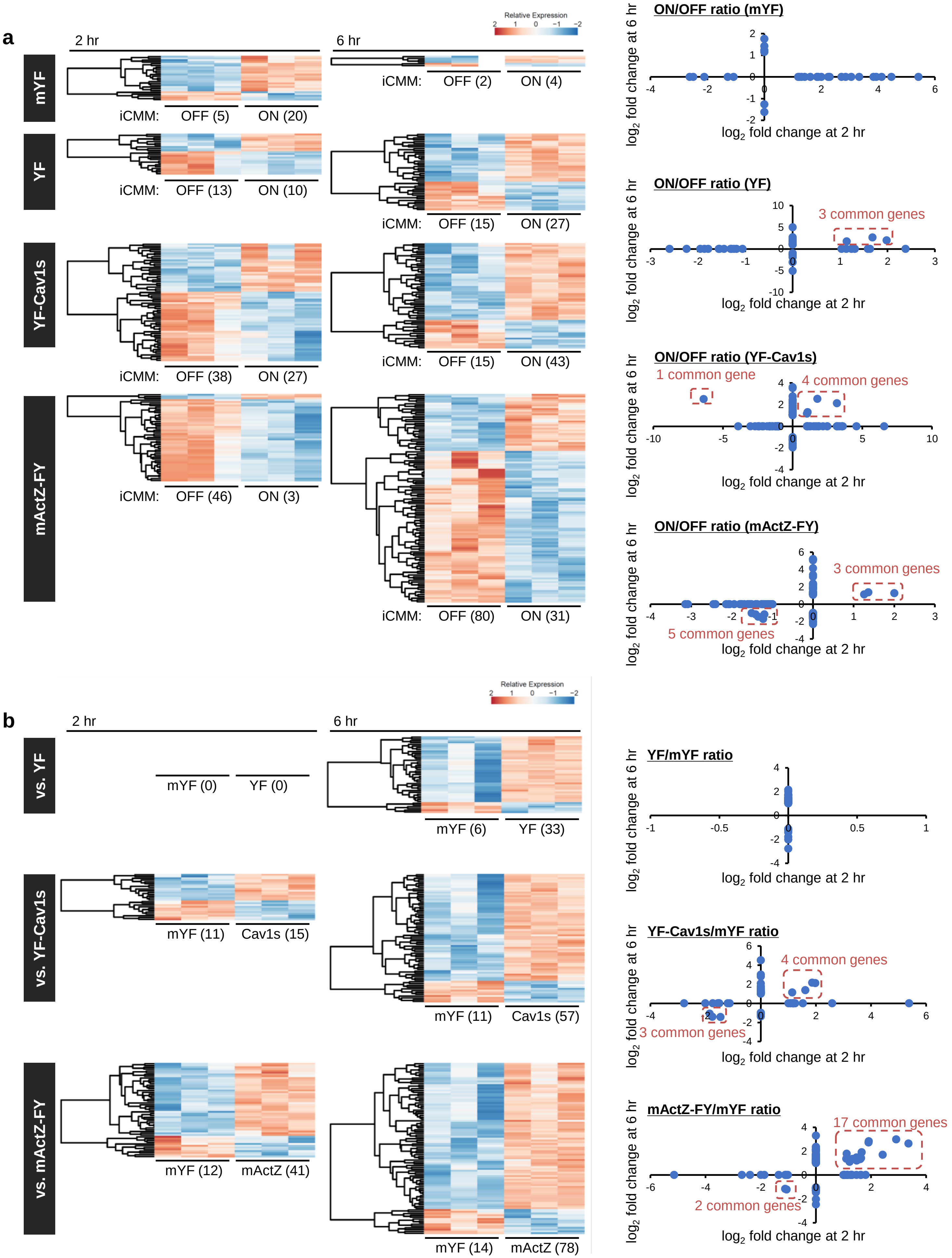


**Supplementary Figure 10. Miyamoto et al.**

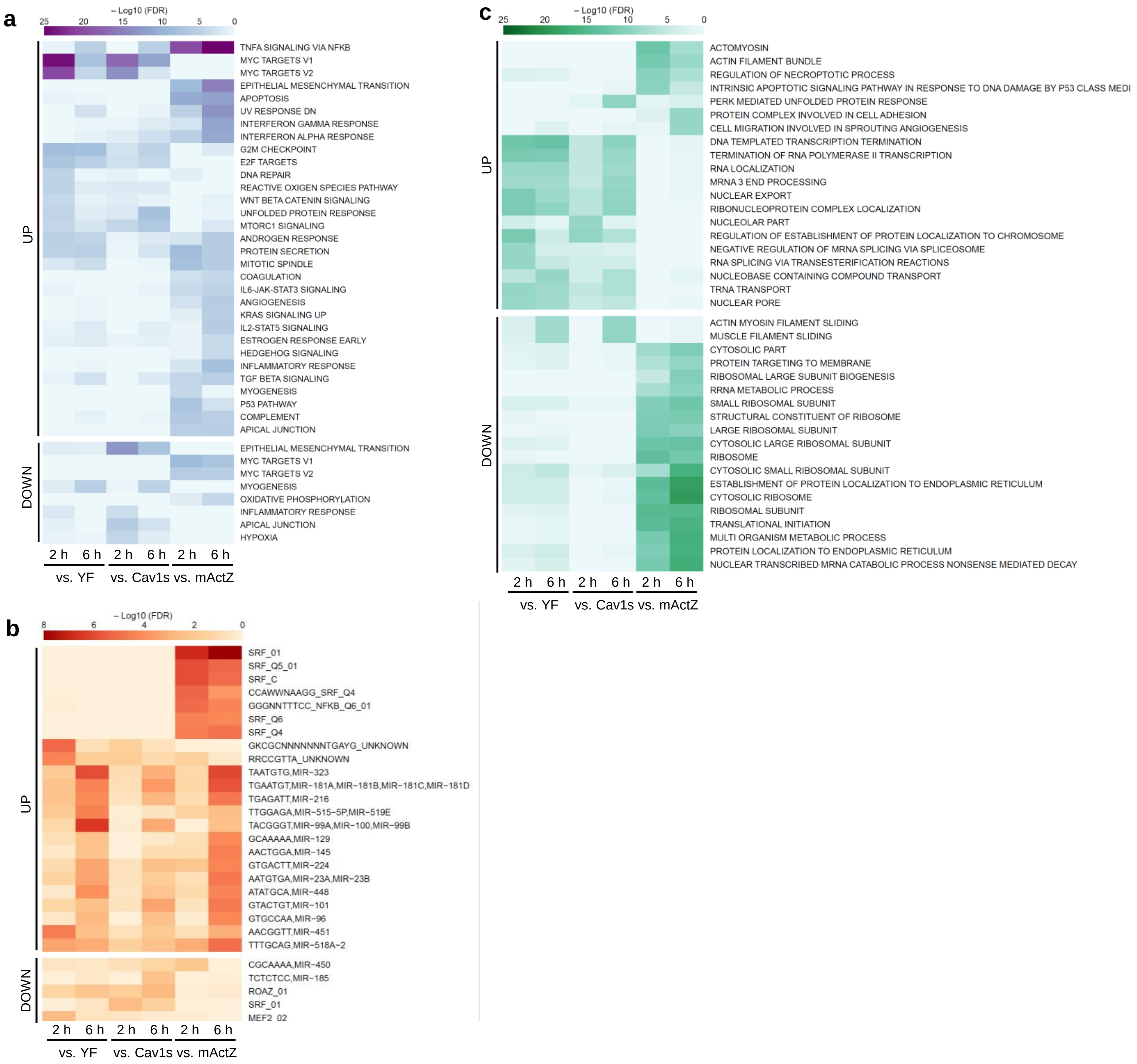


**Supplementary Figure 11. Miyamoto et al.**
